# FeatureMAP enables network-guided interpretation of untargeted metabolomics

**DOI:** 10.64898/2026.11.04.764165

**Authors:** Yuting Wang, Feifan Zhang, Guoping Peng, Lifeng Qiu, Li Zeng, Chuchu Wang, Yijiang Liu, Jiramet Kinchagawat, Xicheng Yang, Chang Liu, Caroline Johnson, Qi Su, Xiaotao Shen

**Affiliations:** Lee Kong Chian School of Medicine, Nanyang Technological University, 59 Nanyang Drive, Singapore 636921; School of Chemistry, Chemical Engineering and Biotechnology, Nanyang Technological University, 62 Nanyang Drive, Singapore 637459; Department of Neurology, The First Affiliated Hospital, Zhejiang University School of Medicine, Hangzhou, China; Neural Stem Cell Research Lab, National Neuroscience Institute, Singapore, Singapore; Neuroscience & Behavioural Disorders Program, Duke-NUS Graduate Medical School, Singapore, Singapore; NTU Institute of Structural Biology, Nanyang Technological University, Singapore, Singapore; State Key Laboratory of Microbial Technology, Shandong University, Qingdao, China; Department of Environmental Health Sciences, Yale School of Public Health, New Haven, CT, USA; Department of Medicine and Therapeutics, State Key Laboratory of Digestive Diseases, Li Ka Shing Institute of Health Sciences, The Chinese University of Hong Kong, Hong Kong, China; Singapore Phenome Center, Nanyang Technological University, 59 Nanyang Drive, Singapore 636921

## Abstract

Untargeted liquid chromatography-mass spectrometry (LC-MS) metabolomics generates thousands of features, but incomplete metabolite annotation and reliance on predefined pathways limit their biological interpretation. Here we present FeatureMAP, an LC-MS feature-centric computational framework that integrates multi-evidence metabolite annotation, network-guided refinement, metabolic module discovery, and structured large language model-based functional interpretation. Across ten public datasets, FeatureMAP improved MS1-based metabolite candidate prioritization while reducing annotation redundancy. In semi-synthetic benchmarks, FeatureMAP recovered coordinated metabolic perturbations, achieving 92% recovery of a simulated perturbation, and distinguished concurrent metabolic processes. Controlled benchmarks further evaluated functional interpretation under competing metabolite-level signals. Based on FeatureMAP, we additionally developed PhenoMAP, a phenotype-aware knowledge graph comprising 12,929 dysregulated metabolic modules from 501 human metabolomics studies spanning 326 phenotypes. Module-level comparisons revealed reproducible metabolic similarities across independent studies and biologically supported relationships among phenotypes. Application to Alzheimer’s disease cerebrospinal fluid and cellular datasets identified convergent membrane-lipid alterations and implicated galactose-linked glycosphingolipid remodeling. Together, FeatureMAP and PhenoMAP enable network-grounded interpretation of incompletely annotated LC-MS metabolomics data and systematic investigation of metabolic dysregulation across biological conditions.

## Introduction

Untargeted liquid chromatography-mass spectrometry (LC-MS) metabolomics provides a powerful approach for profiling small molecules, offering insights into physiological, environmental, and disease-associated biochemical states^1^. However, untargeted LC-MS generates thousands of features defined by mass-to-charge ratio (*m/z*), retention time, and abundance, rather than a matrix of confidently identified metabolites^2^. Translating these extensive feature-level measurements into biologically interpretable metabolic alterations therefore remains a fundamental challenge^2^.

A major obstacle is incomplete metabolite identification. Although tandem mass spectrometry (MS/MS) provides valuable structural evidence, spectral library matching typically annotates only a fraction of detected LC-MS features owing to limited spectral coverage, incomplete reference libraries, and chemical complexity^2,3^. Restricting downstream analyses to confidently identified metabolites consequently excludes much of the acquired information. MS1-based annotation can expand metabolite coverage, but accurate-mass matching frequently yields multiple plausible candidates for a single feature^4^.

Even when metabolites are annotated, their functional interpretation presents another challenge. Conventional pathway enrichment approaches, largely adapted from gene-set analysis, evaluate whether differential metabolites are over-represented in predefined pathways^5^. However, metabolite abundances reflect synthesis, consumption, transport, and exchange rather than pathway activity alone^6^. Consequently, metabolites within the same pathway need not change coordinately^5,7^. Moreover, predefined pathway boundaries may inadequately capture localized or cross-pathway metabolic perturbations, and significance-based feature selection can overlook coordinated but individually modest changes^4,7^.

Several methods have demonstrated the feasibility of interpreting incompletely annotated LC-MS data through biochemical networks^3,4,8,9^. For example, mummichog pioneered the inference of metabolic pathway activity directly from LC-MS features by exploiting the network coherence of plausible metabolite assignments^4^. PIUMet subsequently applied prize-collecting Steiner forest (PCSF) optimization to connect ambiguous features through molecular interaction networks and infer disease-associated subnetworks^8^. More recently, our TidyMass2 framework introduced feature-based functional module analysis within an integrated metabolomics workflow^3^. Despite these advances, important needs remain for integrating complementary annotation evidence with network-guided candidate refinement, identifying robust metabolic modules without predefined significant-feature lists, and systematically interpreting their biological functions while accounting for annotation uncertainty.

Here, we developed FeatureMAP (https://featuremap.jaspershenlab.com/), a feature-centric computational framework integrating multi-evidence metabolite annotation, network-guided module discovery, and structured large language model (LLM)-based functional interpretation. FeatureMAP first integrates *m/z*, retention time, adduct, isotope, and optional MS/MS evidence to generate candidate feature-metabolite assignments. It then incorporates feature-level statistical evidence as continuous weights and applies network refinement to prioritize biochemically coherent candidates. Repeated network optimization, community detection, and statistical evaluation enable the identification of robust dysregulated metabolic modules without requiring prior selection of significant features. Finally, an evidence-guided LLM framework provides structured functional interpretation through complementary fast and multi-agent advanced modes. By integrating all these strategies, FeatureMAP enables biological interpretation beyond individual metabolite identities and predefined pathways.

To extend this approach across studies, we developed PhenoMAP (https://phenomap.jaspershenlab.com/), a phenotype-aware metabolic module atlas integrating FeatureMAP-derived modules, ontology-normalized phenotypes, and metabolites into a knowledge graph. This resource enables module-level comparisons across heterogeneous metabolomics studies. We systematically evaluated FeatureMAP through metabolite annotation benchmarks, simulated metabolic perturbations, and controlled functional interpretation experiments, and assessed cross-study metabolic relationships using PhenoMAP. We further applied FeatureMAP to clinical cerebrospinal fluid (CSF) and cellular models of Alzheimer’s disease (AD) to investigate convergent metabolic alterations. Together, FeatureMAP and PhenoMAP provide a network-guided framework for interpreting incompletely annotated LC-MS data and investigating metabolic dysregulation across biological conditions.

## Results

### Overview of the FeatureMAP framework

FeatureMAP takes an untargeted LC-MS feature table as input and comprises three major stages. First, candidate feature-metabolite assignments are generated using an evidence-integrated annotation strategy adapted from TidyMass2 (Fig. 1a; Supplementary Fig. 1a,b; Methods)^3^. LC-MS features are matched to metabolite databases based on their *m/z* values and grouped into metabolite feature clusters (MFCs) according to complementary ion-level evidence. Iterative evidence scoring and pruning then reduce annotation redundancy while preserving plausible candidate assignments, yielding evidence-integrated MS1-based annotations.

**Fig. 1.**
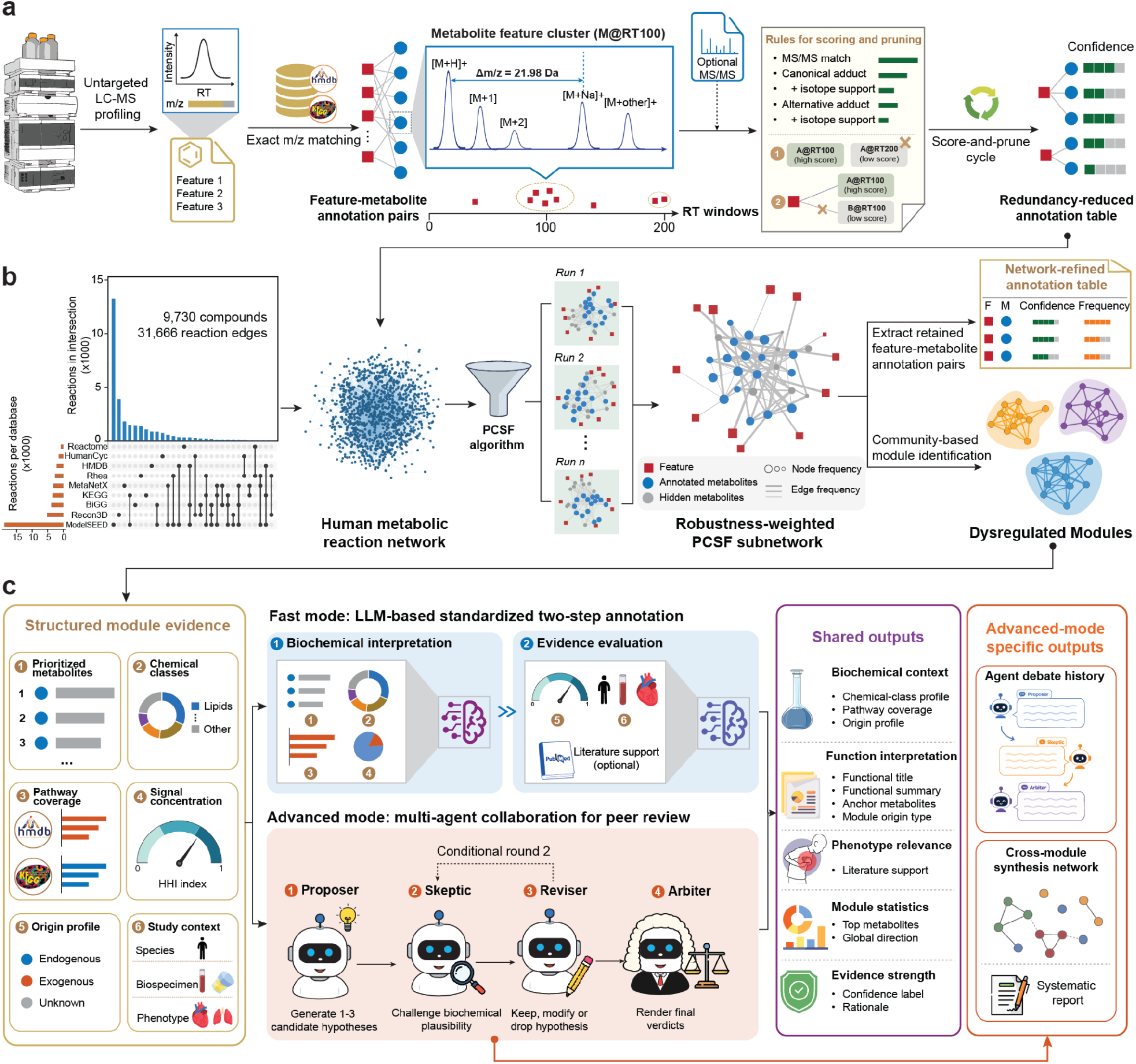
Overview of FeatureMAP. (a) Multi-evidence feature-to-metabolite annotation. LC-MS features are assigned to candidate metabolites by integrating *m/z*, retention time, isotope, adduct, and optional MS/MS evidence, followed by scoring and pruning of redundant assignments. (b) Network-guided discovery of dysregulated metabolic modules. Feature-metabolite assignments and feature-level statistical evidence are 133 integrated with a human metabolic reaction network. Repeated network optimization and community detection generate network-refined annotations and candidate dysregulated metabolic modules. (c) Structured interpretation of dysregulated metabolic modules. Module-level biochemical and study-context evidence is assembled for structured interpretation using either a standardized fast mode or a multi-agent advanced mode.

Second, these candidate annotations are integrated with a human metabolic reaction network to construct a heterogeneous feature-metabolite graph (Fig. 1b; Supplementary Fig. 1c,d; Methods). Feature nodes are assigned continuous weights based on statistical significance and effect size, whereas feature-metabolite edge costs reflect annotation confidence. PCSF optimization identifies biochemically coherent subnetworks while allowing the inclusion of hidden metabolites that connect otherwise disconnected network regions^10^. Repeated optimization under perturbed edge costs generates a robustness-weighted subnetwork, which is further used to refine and rank candidate metabolite annotations. Community detection is then applied to identify metabolic modules, which are evaluated based on network organization, annotation coverage, and feature-level dysregulation to determine statistically significant dysregulated metabolic modules.

Third, the dysregulated metabolic modules are subjected to structured functional interpretation using an evidence-guided LLM framework (Fig. 1c; Methods). Two complementary interpretation modes are provided. The fast mode follows a two-step strategy, first inferring the intrinsic biochemical function of each module and then evaluating its relevance and supporting evidence in the study context. The advanced mode employs a multi-agent review process in which candidate interpretations are proposed, critically evaluated, revised, and adjudicated (Supplementary Fig. 2). Both modes generate structured outputs comprising functional titles and summaries, prioritized anchor metabolites, biochemical context, phenotype relevance, and evidence-strength assessments. The advanced mode additionally preserves the hypothesis development and review history for evidence inspection and optionally integrates accepted interpretations across modules into an interactive synthesis network and study-level report. The methodological differences between FeatureMAP and existing approaches are summarized in Supplementary Table 1.

### FeatureMAP prioritizes MS1-based metabolite candidates through network-guided refinement

Despite the integration of multiple sources of ion-level evidence, individual LC-MS features often remain ambiguously annotated. We hypothesized that correct feature-metabolite assignments are more likely to form coherent metabolic subnetworks than incorrect assignments^4,8^. To evaluate whether network-guided refinement could improve candidate prioritization, we analyzed ten case-control untargeted LC-MS metabolomics datasets (Fig. 2a and Supplementary Fig. 3a; Methods). We evaluated three configurations: initial MS1-based annotations, network-refined annotations, and network-weighted annotations (Methods).

We first evaluated candidate ranking using tie-adjusted annotation accuracy, which accounts for multiple candidates sharing the same rank (Supplementary Methods). Network refinement significantly improved top-1 tie-adjusted annotation accuracy relative to the initial MS1-based annotations, and network-weighted ranking provided a further improvement (Fig. 2b). Similar improvements were observed for the top two and three candidate ranks (Supplementary Fig. 3b). Consistent with these findings, MS/MS-supported reference assignments exhibited significantly higher edge-retention frequencies than alternative candidate assignments across repeated PCSF optimizations (Fig. 2c), indicating that network-derived robustness provides complementary evidence for candidate prioritization.

Network refinement also substantially reduced annotation redundancy, measured as the mean number of candidate metabolites per feature (Fig. 2d and Supplementary Fig. 3c). Among initial annotations mappable to the metabolic reaction network, MS/MS-supported reference assignments were retained at a significantly higher rate than alternative candidates (Supplementary Fig. 3d). Most retained reference assignments ranked among the top three candidates (Fig. 2d). For example, feature M407T446_NEG matched cholic acid by MS/MS spectral matching, whereas its initial MS1-based annotations included numerous bile-acid candidates with similar scores. Network-weighted ranking placed cholic acid above these alternatives (Fig. 2e; Supplementary Data 1).

We next compared FeatureMAP with mWISE and xMSannotator using MS/MS-supported features evaluable by all three methods (Methods)^11,12^. FeatureMAP achieved significantly higher top-1 tie-adjusted annotation accuracy and F1 scores than both comparators (Fig. 2f). Its annotation redundancy was comparable to mWISE and substantially lower than xMSannotator (Fig. 2g and Supplementary Fig. 3e). Together, these results demonstrate that metabolic network optimization improves the retention and prioritization of MS/MS-supported metabolite candidates while reducing annotation redundancy, providing a refined annotation basis for downstream dysregulated metabolic module discovery.

### FeatureMAP recovers metabolic perturbations through network-guided module discovery

We next evaluated whether FeatureMAP could identify coordinated metabolic perturbations from LC-MS feature-level data. We constructed a semi-synthetic benchmark using untargeted metabolomics data from 40 control participants from a gestational diabetes cohort, randomly divided into pseudo-case and pseudo-control groups (n = 20 each)^13^. As expected, no FDR-significant differential features or dysregulated metabolic modules were detected between the groups before perturbation (Supplementary Fig. 4a and Methods). We selected arginine biosynthesis as the target pathway, with seven of its metabolites represented by 11 LC-MS features supported by MS/MS spectral matching (Fig. 3a, Supplementary Fig. 4b and Supplementary Data 2). Across 100 simulation runs, these features were randomly increased 2- to 10-fold in the pseudo-case group (Methods). Directly perturbed metabolites were defined as spiked metabolites, whereas other metabolites in the same pathway that were not directly modified were defined as unspiked metabolites. A simulation was considered successful if exactly one significant dysregulated metabolic module was identified and contained at least five of the seven spiked metabolites.

**Fig. 2.**
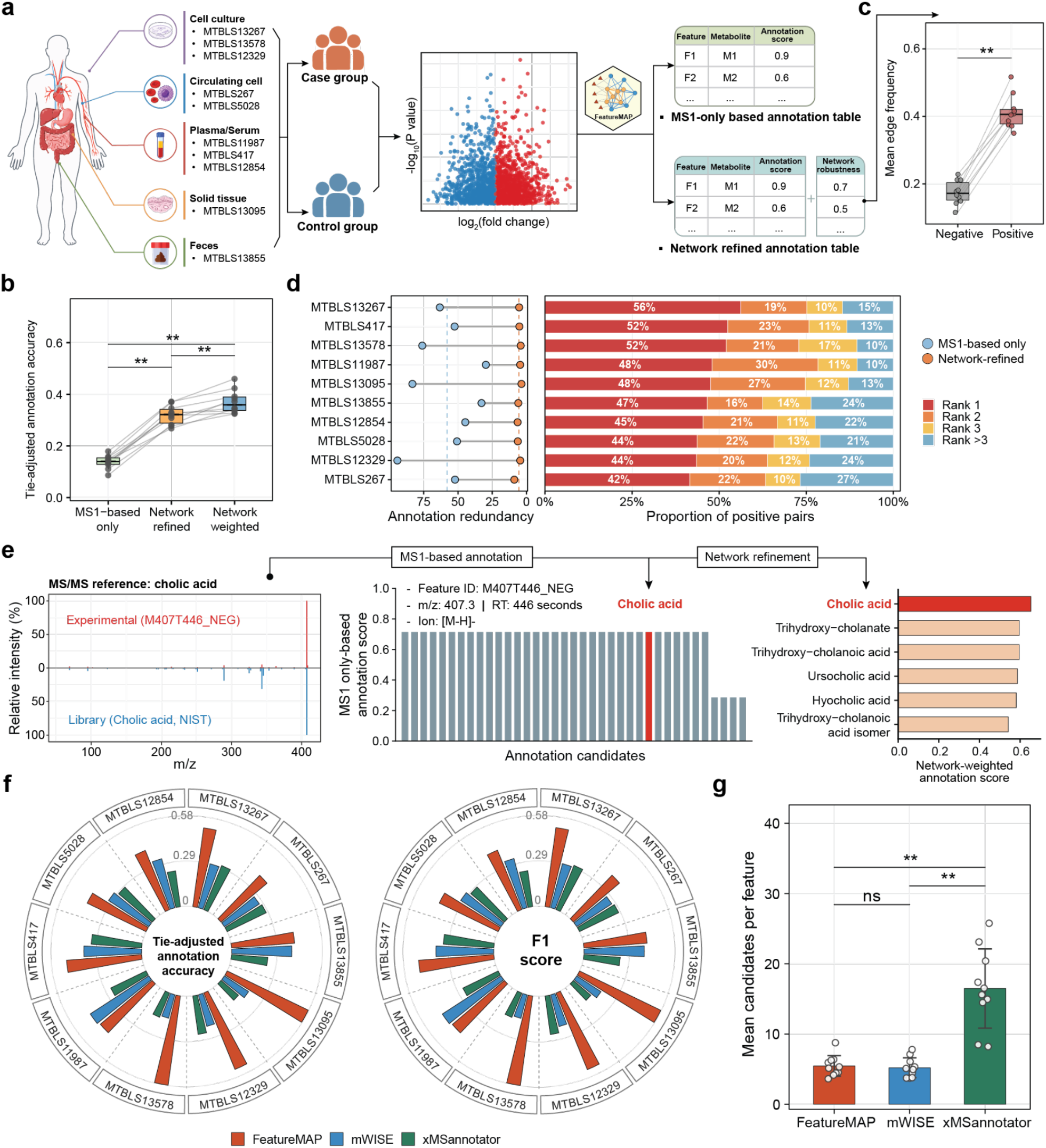
FeatureMAP prioritizes MS1-based metabolite candidates through network-guided refinement. (a) Benchmark design using ten MetaboLights datasets spanning diverse biological sample types. (b) Tie-adjusted annotation accuracy before and after network refinement and network weighting. (c) Mean edge frequency of positive and negative feature-metabolite assignments across perturbed PCSF optimizations. Positive assignments are feature-metabolite pairs matching the MS/MS-supported reference annotation. Negative assignments are alternative candidates for the same features. (d) Annotation redundancy (left) and rank distribution of positive assignments after network refinement (right). (e) Network-guided prioritization of the MS/MS-supported cholic acid annotation for feature M407T446_NEG. (f) Comparison of top-rank tie-adjusted annotation accuracy and F1 score between FeatureMAP, mWISE, and xMSannotator. (g) Mean number of candidate metabolites reported per feature by each method. Points represent individual datasets (n = 10). Statistical significance was assessed using two-sided paired Wilcoxon signed-rank tests. ns, not significant; **P < 0.01.

**Fig. 3.**
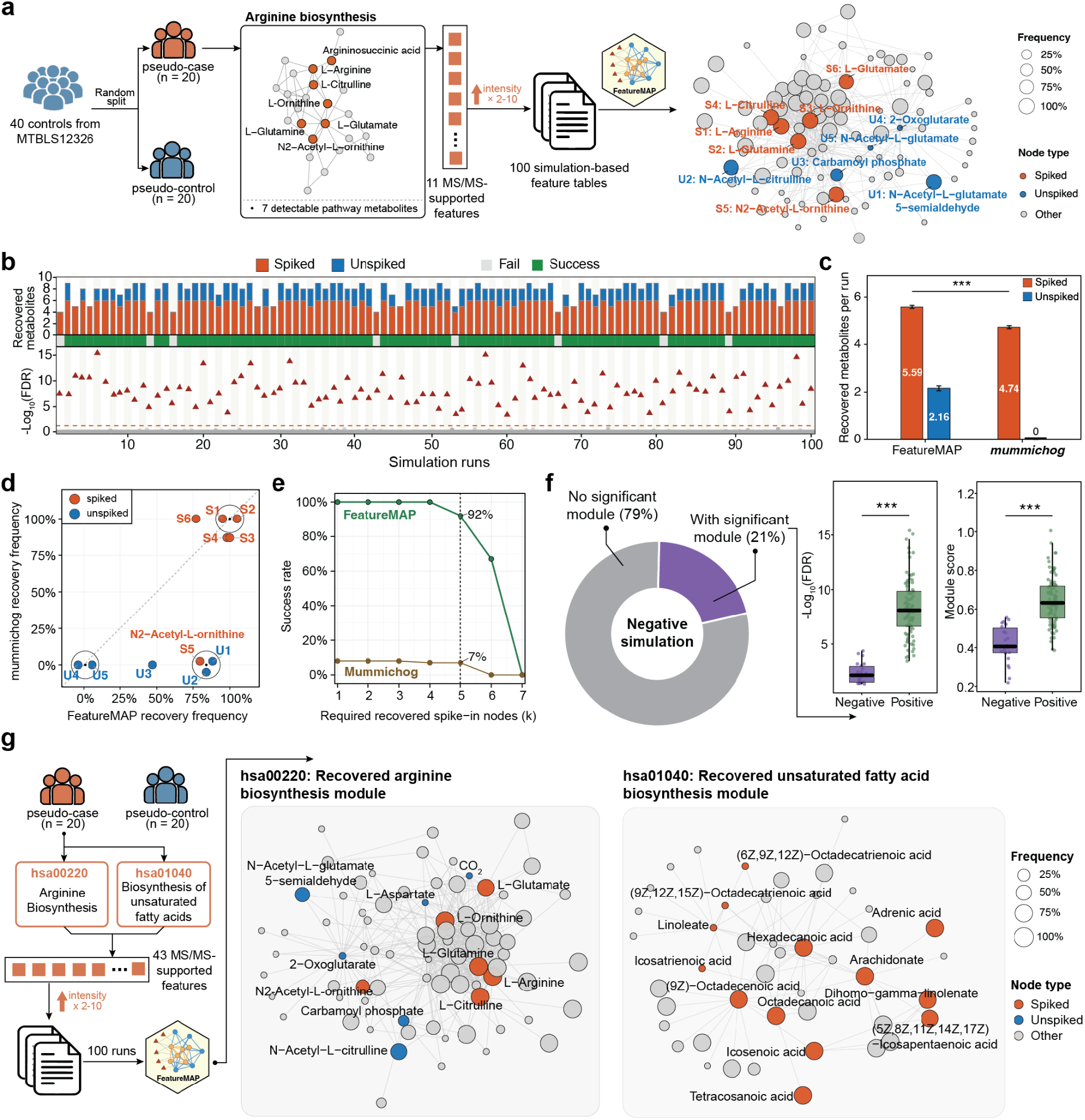
FeatureMAP recovers simulated metabolic perturbations as dysregulated metabolic modules. (a) Simulation design using 40 control samples divided into pseudo-case and pseudo-control groups. Eleven MS/MS supported features representing seven metabolites in arginine biosynthesis were increased 2- to 10-fold across 100 simulation runs. (b) Recovery of spiked and unspiked metabolites and statistical significance of the recovered modules across individual runs. (c) Mean numbers of spiked and unspiked metabolites recovered by FeatureMAP and mummichog. Data are presented as the mean ± s.e.m. across 100 simulation runs. Statistical significance was assessed using a two-sided paired Wilcoxon signed-rank test; ***: P < 0.001. (d) Recovery frequencies of individual pathway metabolites by the two methods. Circled points represent metabolites with identical frequencies; labels correspond to those in (a). (e) Success rates under increasingly stringent recovery criteria. A simulation was considered successful only when exactly one significant module was identified, and that module contained at least k spiked metabolites. The vertical dashed line indicates the predefined threshold of k = 5. (f) Frequency of significant modules in negative-control simulations and comparison of their statistical significance and module scores with pathway-spike simulations. Statistical significance was assessed using two sided Wilcoxon rank-sum tests; ***P < 0.001. (g) Concurrent perturbation and recovery of distinct modules representing arginine biosynthesis and unsaturated fatty acid biosynthesis.

FeatureMAP identified exactly one significant module in all 100 simulations, with the resulting modules containing both directly perturbed metabolites and additional unspiked pathway metabolites (Fig. 3b). Under the predefined success criterion, 92% of simulations successfully recovered the target perturbation, with similar results obtained using alternative community-detection algorithms (Supplementary Fig. 4c). To benchmark module recovery, we applied mummichog to the same simulated datasets. FeatureMAP identified an average of 5.59 spiked metabolites per run, compared with 4.74 for mummichog, and additionally included an average of 2.16 unspiked metabolites, none of which were recovered in the selected mummichog modules (Fig. 3c). Metabolite-level analysis further showed that FeatureMAP more consistently recovered N2-acetylornithine and several unspiked pathway intermediates (Fig. 3d). These findings indicate that FeatureMAP captures not only directly perturbed metabolites but also their surrounding biochemical network context.

We further assessed the robustness of these findings by varying the minimum number of spiked metabolites required for successful module recovery while maintaining the requirement of exactly one significant module. FeatureMAP consistently achieved higher success rates than mummichog across thresholds requiring one to six spiked metabolites, including 92% versus 7% at the predefined threshold of five metabolites (Fig. 3e). Thus, the comparative advantage of FeatureMAP was maintained across a range of recovery criteria.

To examine whether module discovery preferentially captured coordinated rather than randomly distributed metabolic alterations, we performed negative-control simulations in which seven randomly selected MS/MS-supported metabolites were perturbed using the same experimental background and fold-change range (Methods). FeatureMAP identified no significant modules in 79 of 100 simulations, whereas the remaining 21 yielded at least one significant module (Fig. 3f). However, these modules showed limited overlap in metabolite composition across simulations (Supplementary Fig. 4d) and had significantly lower −log_10_(FDR) values and module scores than those identified in the arginine-biosynthesis simulations (Fig. 3f). These results suggest that coordinated metabolic perturbations generate stronger and more consistent module-level evidence than randomly distributed changes.

We further investigated whether FeatureMAP could resolve multiple metabolic processes perturbed simultaneously. We extended the benchmark to include concurrent perturbations of arginine biosynthesis and unsaturated fatty-acid biosynthesis, involving 25 spiked metabolites (Fig. 3g, Supplementary Data 2 and Methods). FeatureMAP identified distinct modules corresponding to the two pathways across 100 simulations, with both modules reaching FDR significance in 82% of runs (Supplementary Fig. 4e). Representative subnetworks confirmed that the two perturbations were resolved into separate modules rather than merged into a single network (Fig. 3g). Together, these benchmarks demonstrate that FeatureMAP can identify coordinated metabolic perturbations, capture their connected biochemical neighborhoods, and distinguish concurrent metabolic processes within the same dataset.

### FeatureMAP enables noise-aware functional interpretation of dysregulated metabolic modules

We next evaluated whether FeatureMAP could accurately interpret metabolic modules in the presence of competing biochemical signals. To establish a controlled benchmark, we constructed 208 semi-synthetic modules from 104 curated human PathBank pathways, each comprising reference pathway metabolites, neighboring metabolites, and noise metabolites (Fig. 4a and Supplementary Data 3; Methods)^14^. For each pathway, we generated two modules: a signal-dominant (easy) module, in which reference metabolites received high importance and robustness weights while noise metabolites received low weights, and a noise-confounded (hard) module, in which noise metabolites received weights comparable to those of reference metabolites. Neighboring metabolites received intermediate weights in both settings.

**Fig. 4.**
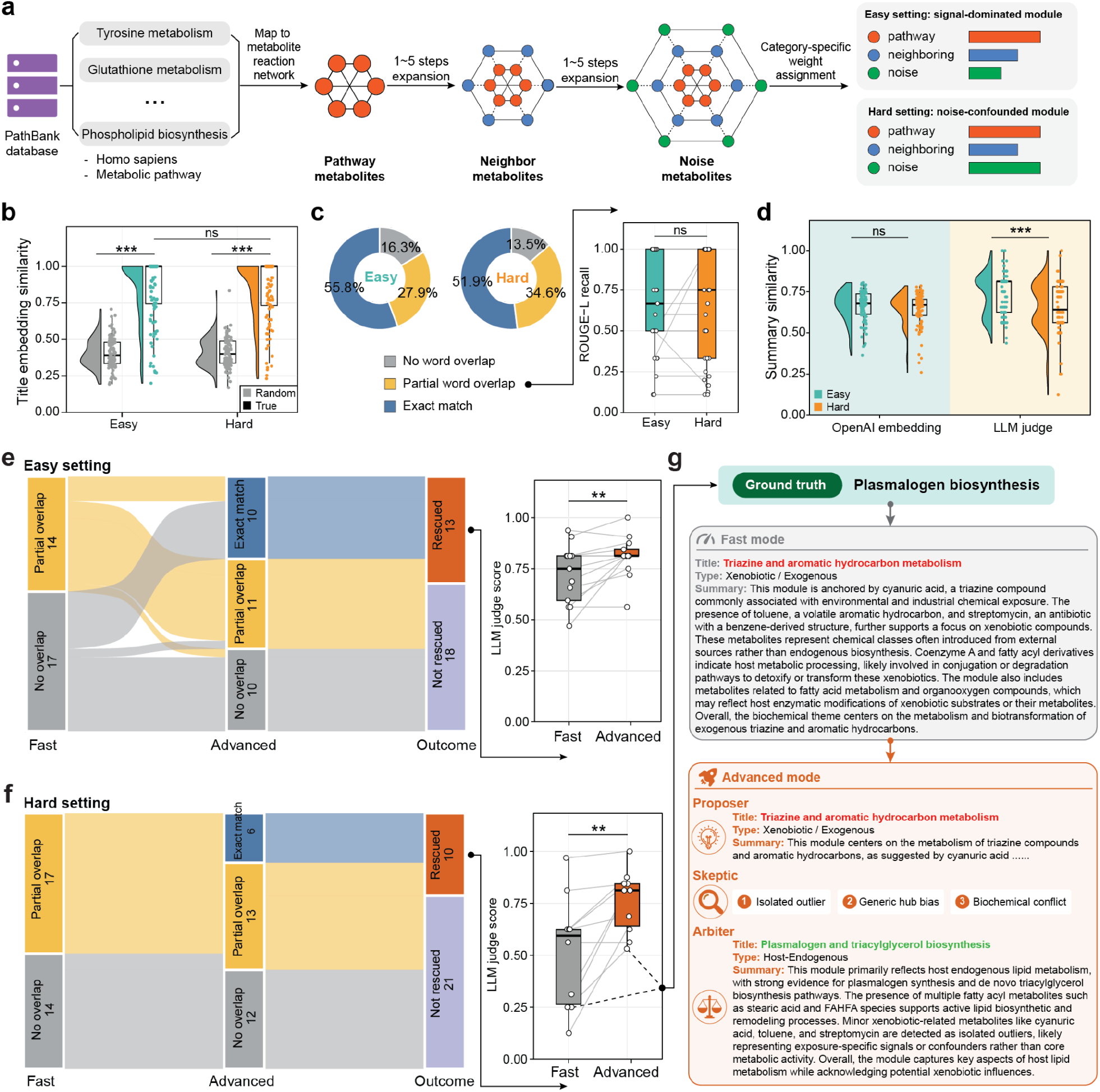
FeatureMAP enables noise-aware functional interpretation of metabolic modules. (a) Construction 13 of paired easy and hard benchmark modules by combining pathway metabolites with neighboring and noise metabolites sampled from the metabolic reaction network and assigned different weights. (b) Semantic similarity of generated titles to their corresponding reference titles or randomly paired titles (n = 104 modules per condition). Two-sided Wilcoxon signed-rank tests were used. ***P < 0.001; ns, not significant. (c) Exact, partial, and absent word overlap between generated and reference titles. ROUGE-L recall for partial matches was compared between easy and hard settings using a two-sided Wilcoxon rank-sum test; ns, not significant. (d) Agreement of generated summaries with the reference pathway descriptions, assessed by embedding similarity and LLM-judge scores. Easy and hard settings were compared using two-sided Wilcoxon signed-rank tests. (e) Advanced-mode reanalysis of weak fast-mode interpretations in the easy setting. The Sankey diagram shows changes in title overlap and rescue status. Paired LLM-judge scores are shown for the 13 rescued modules. (f) Advanced-mode reanalysis in the hard setting. Paired LLM-judge scores are shown for the 10 rescued modules. (g) Representative rescue of plasmalogen biosynthesis through proposer-skeptic-arbiter reasoning in advanced mode.

We first assessed whether FeatureMAP fast mode could recover the underlying pathway identities from these modules. In both settings, generated functional titles showed significantly higher embedding similarity to their corresponding reference pathway names than to randomly paired names (Fig. 4b and Supplementary Fig. 5a). Exact title matches were achieved for 55.8% of easy modules and 51.9% of hard modules, with partial overlap in an additional 27.9% and 34.6%, respectively (Fig. 4c; Methods). Neither title embedding similarity nor ROUGE-L recall among partially matched titles differed significantly between settings (Fig. 4b,c)^15^. These findings suggest that fast mode can recover the principal biochemical themes even when noise metabolites carry substantial weights.

We next examined whether this robustness extended to more detailed functional summaries. Although embedding similarity between generated summaries and reference pathway descriptions was comparable across settings, hard modules received significantly lower functional compatibility scores from a forward LLM judge evaluating thematic agreement, metabolite support, mechanistic consistency, and unsupported claims (Fig. 4d and Supplementary Table 2; Methods). Complementary evaluations using MedCPT embeddings and a reverse judge assessing interpretation errors supported these findings (Supplementary Fig. 5b-d and Supplementary Table 3). Forward compatibility and reverse error scores were strongly inversely correlated in both the easy and hard settings (Spearman ρ = −0.822 and −0.820, respectively; both P < 0.001; Supplementary Fig. 5e,f). Together, these results indicate that, although fast mode generally preserves pathway-level semantic information, detailed functional interpretations are more sensitive to competing metabolite signals.

To determine whether more extensive reasoning could improve difficult interpretations, we reanalyzed modules with weak fast-mode title recovery using advanced mode. Modules with no title overlap or partial overlap with below-median ROUGE-L recall were selected, yielding 31 easy and 31 hard modules. Advanced mode improved title recovery in 13 easy modules and 10 hard modules by converting previously unmatched titles into partial or exact matches or increasing ROUGE-L recall for partially matched titles (Fig. 4e,f; Supplementary Data 4 and 5). Among these rescued modules, functional compatibility scores also increased significantly, indicating that multi-agent refinement improved both title recovery and summary-level agreement with the reference descriptions in a subset of challenging cases.

A representative plasmalogen-biosynthesis module illustrates this improvement (Fig. 4g). Fast mode assigned the title “Triazine and aromatic hydrocarbon metabolism”, reflecting excessive emphasis on xenobiotic-related noise metabolites. During advanced-mode interpretation, the Proposer initially proposed this hypothesis, but the Skeptic identified isolated-outlier bias, generic-hub bias, and biochemical inconsistencies. Following critique and revision, the Arbiter accepted “Plasmalogen and triacylglycerol biosynthesis”, restoring the reference pathway theme within a broader lipid-metabolism interpretation. Together, these benchmarks demonstrate that FeatureMAP fast mode can provide consistent pathway-level interpretations, whereas advanced mode offers additional refinement for selected modules with ambiguous or noise-confounded functional annotations.

### PhenoMAP reveals reproducible metabolic relationships across human phenotypes

We next investigated whether FeatureMAP-derived modules could enable comparisons across heterogeneous human metabolomics studies. To this end, we systematically analyzed publicly available untargeted LC-MS datasets using a standardized TidyMass-FeatureMAP workflow (Fig. 5a and Methods)^16,17^. We retained 501 studies yielding at least one significant dysregulated metabolic module and integrated their results into PhenoMAP, a phenotype-aware metabolic module atlas portal.

**Fig. 5.**
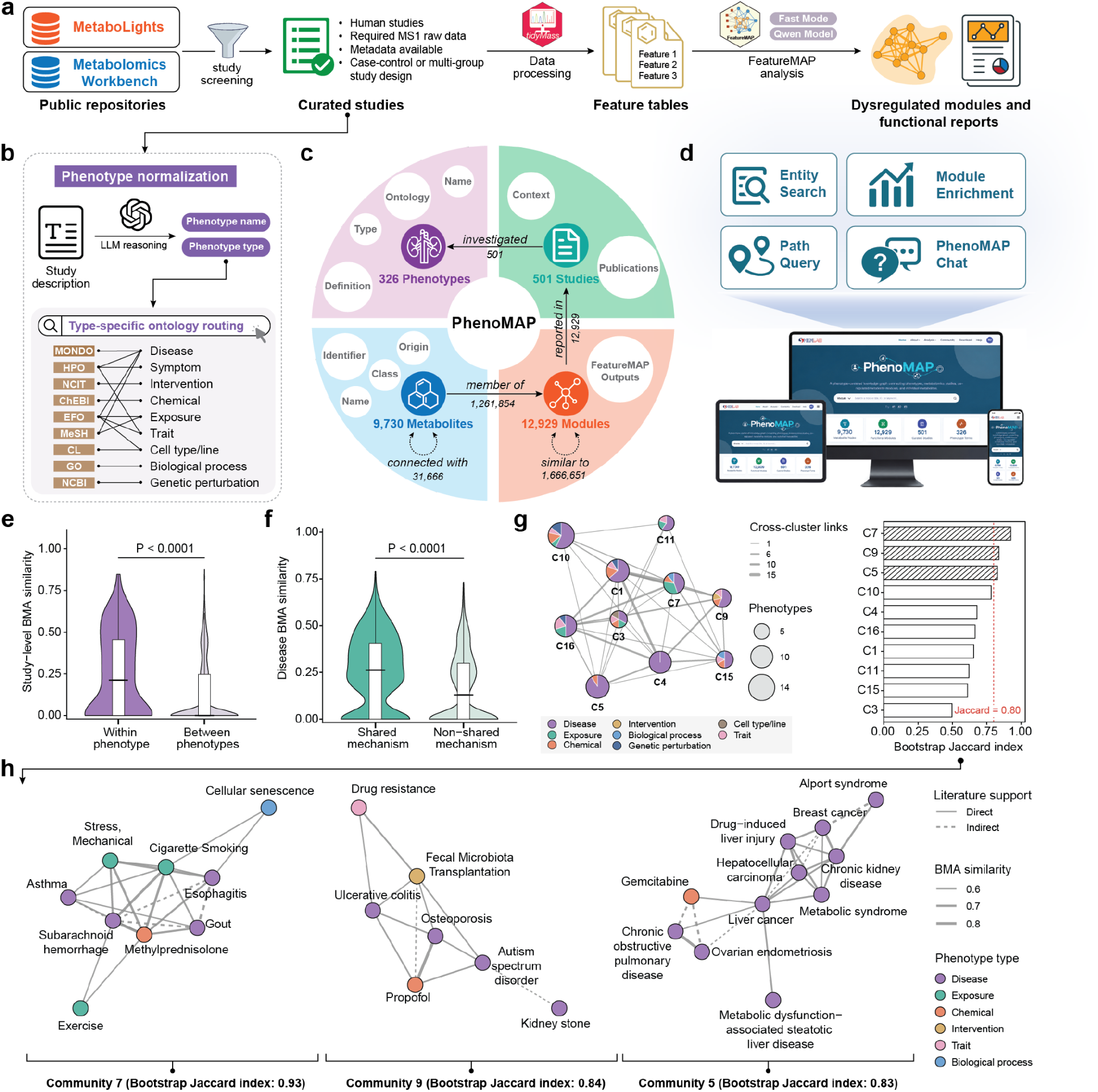
PhenoMAP enables phenotype-aware integration and cross-study comparison of dysregulated metabolic modules. (a) Overview of study curation and module generation for PhenoMAP. (b) Phenotype normalization and type-specific ontology mapping. (c) Entities and relationships in the PhenoMAP knowledge graph. (d) PhenoMAP interfaces for entity search, path query, module enrichment, and natural-language chat. (e) Best-match-average similarity between study-level module sets from the same phenotype or different phenotypes. (f) Best-match-average similarity between disease pairs with or without a shared metabolic mechanism, defined using CTD-curated KEGG pathway associations. (g) Consensus communities in the phenotype-similarity network and their bootstrap stability. Node size indicates the number of phenotypes, sectors indicate phenotype-type composition, and edge width indicates the number of cross-community links. The dashed line marks a bootstrap Jaccard index of 0.80. (h) Representative phenotype communities. Node color indicates phenotype type, edge width indicates best-match-average similarity, and solid and dashed edges indicate direct and indirect literature support, respectively.

To enable phenotype-level comparisons, we developed an LLM-assisted workflow to extract phenotypes from study descriptions and map them to type-specific biomedical ontologies, resulting in 326 standardized phenotype entities across nine categories (Fig. 5b). We then constructed a knowledge graph linking these phenotypes and their associated studies to 12,929 dysregulated metabolic modules and 9,730 metabolites through relationships representing study context, module provenance, metabolite membership, biochemical reactions and module similarity (Fig. 5c). To facilitate exploration of these relationships, we developed an interactive web portal supporting entity search, graph-based path queries, module enrichment analysis of external metabolite lists and graph-grounded natural-language queries (Fig. 5d). These functionalities were illustrated using purine metabolism as a representative example (Supplementary Fig. 6 and Supplementary Note 1).

We first evaluated whether independently generated studies investigating the same phenotype exhibited greater module-level similarity than studies investigating different phenotypes. Using a bidirectional best-match average (BMA) to compare study-level module sets, we identified 122 same-phenotype and 4,064 different-phenotype study pairs across 31 phenotypes represented in at least two independent studies (Methods)^18^. Same-phenotype study pairs exhibited significantly higher BMA similarity than different-phenotype pairs (P < 0.0001, one-sided Wilcoxon rank-sum test; Fig. 5e), supporting the reproducibility of phenotype-associated metabolic module patterns across independent studies. We next examined whether module-based similarities between distinct diseases were consistent with established metabolic knowledge. Using Comparative Toxicogenomics Database (CTD)-curated KEGG pathway associations for 61 diseases, we found that disease pairs with above-average numbers of shared pathway associations exhibited significantly higher BMA similarity than the remaining pairs (median, 0.262 versus 0.130; P < 0.0001, one-sided Wilcoxon rank-sum test; Fig. 5f; Methods). These findings indicate that module-level similarities reflect both recurring metabolic patterns across studies of the same phenotype and relationships consistent with curated metabolic-pathway knowledge across different diseases.

We further investigated whether metabolic similarities could organize heterogeneous phenotypes into biologically supported communities. Consensus community detection identified communities spanning diseases, exposures, interventions, traits, and biological processes, with C7, C9, and C5 showing particularly high stability (Fig. 5g,h and Supplementary Fig. 7). C7 connected cigarette smoking, asthma, cellular senescence, and methylprednisolone, with arachidonic-acid-related modules recurring across these phenotypes. C9 included ulcerative colitis and fecal microbiota transplantation, capturing a documented disease-intervention relationship. C5 grouped renal, hepatic and metabolic diseases, including Alport syndrome, chronic kidney disease, metabolic syndrome, metabolic dysfunction-associated steatotic liver disease and liver cancer, with representative module correspondences involving sphingolipid, fatty-acid and glycerophospholipid metabolism. Literature curation supported most displayed relationships directly and the remainder through shared biological mechanisms (Methods; Supplementary Data 6–8).

Together, PhenoMAP provides a unified resource for integrating dysregulated metabolic modules across human metabolomics studies. The observed cross-study consistency, concordance with curated metabolic-pathway knowledge, and recovery of biologically supported phenotype communities demonstrate its potential for investigating shared metabolic dysregulation across diverse human conditions.

### FeatureMAP reveals module-level metabolic convergence across clinical and cellular models of AD

Finally, we evaluated whether FeatureMAP could identify convergent metabolic modules across clinical and cellular systems. We independently analyzed CSF samples from 21 patients with AD and 21 age- and sex-matched individuals with other neurological or medical conditions, together with AD-derived and control induced pluripotent stem cell (iPSC) samples (n = 6 per group; Fig. 6a and Supplementary Fig. 8a,b). Nominal feature-level differences were widespread: 970 of 16,240 hydrophilic interaction liquid chromatography (HILIC) and 1,097 of 20,923 reversed-phase chromatography (RP) features differed in CSF, compared with 347 of 13,982 HILIC and 850 of 14,153 RP features in iPSCs (P < 0.05; Fig. 6b). This diffuse signal was not resolved by conventional metabolite-centered analysis. Of the 2,146 and 1,597 Level 1/2-annotated metabolites in CSF and iPSCs, respectively, 168 and 94 were nominally differential, but none remained significant after multiple-testing correction. Conventional KEGG enrichment likewise identified no nominally significant pathway in either dataset (Supplementary Fig. 8c,d).

**Fig. 6.**
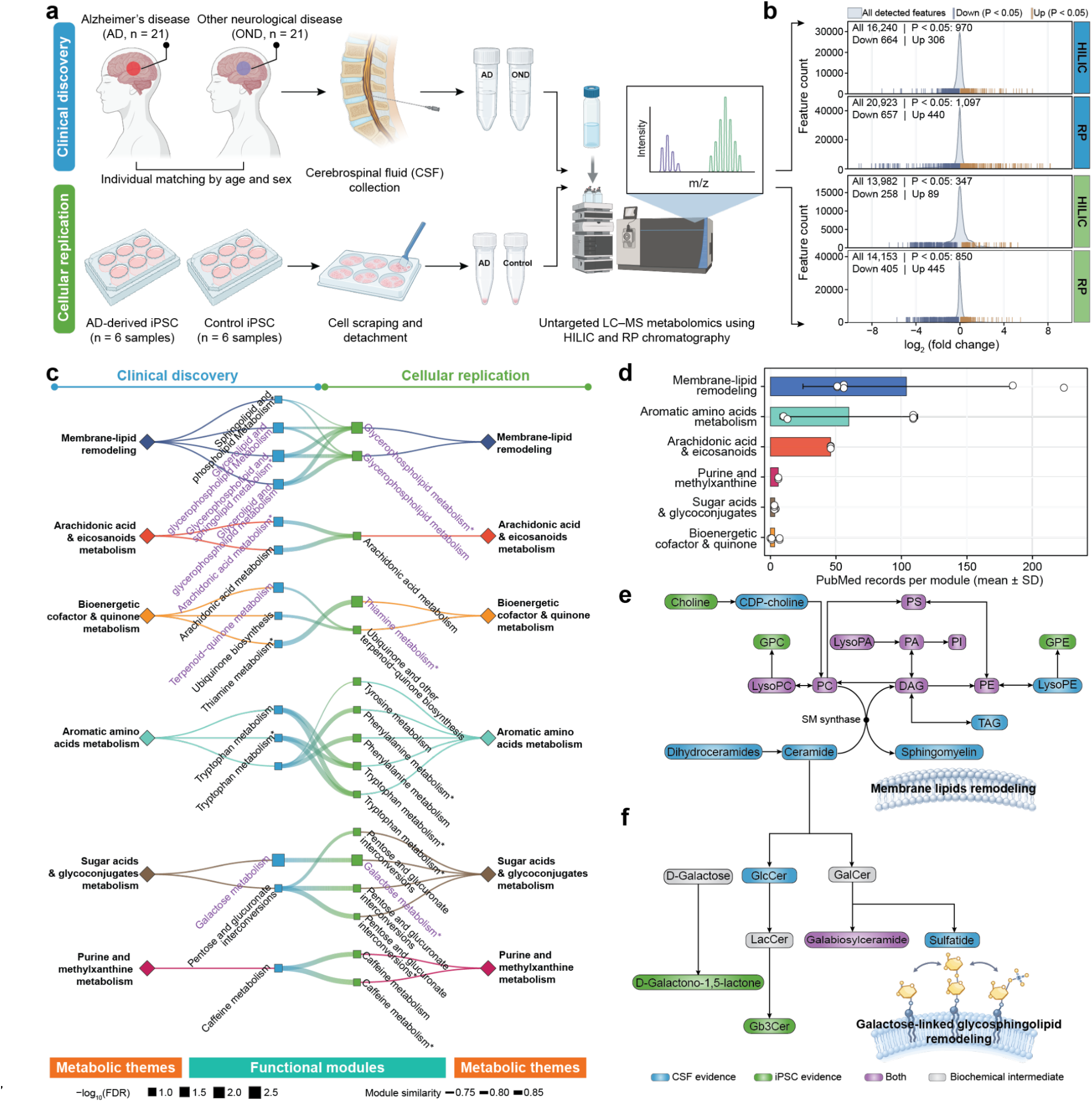
FeatureMAP reveals module-level metabolic convergence across clinical and cellular models of AD. (a) Study design for clinical discovery in CSF and AD-derived iPSC models. (b) Distributions of feature-level log_2_(fold change) across HILIC and RP modes. Colored rug marks denote nominally differential features (P < 0.05) and their directions. (c) Correspondence between CSF and iPSC modules grouped by metabolic theme. Diamonds represent metabolic themes and squares represent modules. Square size indicates −log_10_(FDR), edge width indicates module similarity, and purple module labels denote modules with FDR < 0.05. Only module pairs with similarity ≥ 0.70 are shown. Asterisks denote HILIC-derived modules. (d) AD PubMed records retrieved for modules in each metabolic theme. Bars and error bars show mean ± s.d., and points represent individual modules. (e) Metabolite map of membrane-lipid remodeling integrating evidence from the CSF and iPSC analyses. (f) Metabolite map linking galactose metabolism to glycosphingolipid remodeling in the CSF and iPSC analyses. In (e) and (f), blue, green, and purple nodes indicate CSF, iPSC, and shared evidence, respectively. Gray nodes indicate connecting biochemical intermediates. Arrows represent known biochemical relationships.

FeatureMAP instead integrated evidence from all detected features, rather than a significance-filtered list of annotated metabolites. Network refinement improved annotation accuracy against Level 1 references and reduced candidate redundancy in both datasets (Supplementary Fig. 8e). Subsequent module analysis identified 30 nominally significant, high-confidence host-endogenous modules in CSF and 40 in iPSCs (Supplementary Fig. 8f,g). Fifteen CSF and 16 iPSC modules formed 26 cross-system correspondences with similarity ≥ 0.70 (Fig. 6c; Supplementary Data 9 and 10). These correspondences organized into six coherent themes spanning membrane-lipid remodeling, arachidonic acid and eicosanoid metabolism, bioenergetic cofactor and quinone metabolism, aromatic amino-acid metabolism, sugar-acid and glycoconjugate metabolism, and purine and methylxanthine metabolism. Five of the six membrane-lipid modules and both galactose-metabolism modules remained significant after module-level FDR correction.

Literature support further distinguished an established from a less-characterized component of this convergence. Membrane-lipid remodeling retrieved the largest number of AD-related PubMed records and therefore provided a biologically established benchmark (Fig. 6d). Anchors from four CSF and two iPSC modules reconstructed a common network linking phosphatidylcholine turnover, glycerophospholipid remodeling, glycerolipid metabolism, and ceramide–sphingomyelin metabolism (Fig. 6e and Supplementary Data 11). The two systems converged on a central LysoPA–PA–PI–PS– PC–LysoPC–PE–DAG network, while extending into complementary branches: CSF captured CDP-choline, LysoPE, triacylglycerols, ceramides, and sphingomyelins, whereas iPSCs captured choline, glycerophosphocholine, and glycerophosphoethanolamine. Level 1/2 evidence independently supported key sphingolipid anchors in CSF (Cer(d18:0/16:0), Cer(d18:0/18:0)) and glycerophospholipid anchors in iPSCs (glycerophosphocholine, glycerophosphoethanolamine, LysoPC(16:0), LysoPC(16:1), and PC(16:1/16:1)). For each directly matched feature, the corresponding metabolite was ranked first among candidate annotations after network refinement in FeatureMAP (Supplementary Fig. 9).

Against this established lipid background, galactose metabolism represented a comparatively underexplored but equally reproducible signal. The corresponding CSF and iPSC modules were both FDR significant and showed high similarity (module-module similarity = 0.855), despite the limited AD literature associated with this theme. CSF prioritized glucosylceramide and sulfatide, whereas iPSCs prioritized D-galactono-1,5-lactone and globotriaosylceramide (Gb3) (Fig. 6f and Supplementary Data 12). Notably, galabiosylceramide (Gb2) was independently prioritized in both datasets. Curated mapping placed these candidates within complementary galactose/sugar-acid and glycosphingolipid branches. Ceramide anchors from the membrane-lipid modules linked these glycosphingolipid candidates to the broader membrane-remodeling network. This cross-system evidence nominates galactose-linked, ceramide-dependent glycosphingolipid remodeling as a testable component of AD-associated metabolic dysregulation, while retaining the need for targeted confirmation of several molecular identities.

Additional cross-system correspondences involved thiamine metabolism (similarity = 0.863), ubiquinone and terpenoid-quinone metabolism (0.754-0.779), and pentose and glucuronate interconversions (0.793-0.869; Supplementary Fig. 8g; Supplementary Data 9 and 10). Together, these results demonstrate that FeatureMAP enables module-level comparisons across clinical and cellular metabolomics datasets, recovering established lipid-associated alterations while highlighting additional metabolic processes for targeted investigation.

## Discussion

Here, we developed FeatureMAP, a feature-centric computational framework that bridges the gap between incompletely annotated LC-MS features and biologically interpretable metabolic dysregulation. By integrating multi-evidence annotation, feature-level statistical information, and metabolic network topology, FeatureMAP enables network-guided metabolite candidate prioritization, robust discovery of dysregulated metabolic modules and structured LLM-based functional interpretation without requiring complete metabolite identification. Systematic benchmarks demonstrated improvements in annotation refinement, recovery of coordinated metabolic perturbations, and functional interpretation under competing biochemical signals. We further extended FeatureMAP into PhenoMAP, a phenotype-aware metabolic module atlas, and demonstrated its potential for cross-study comparison and biological hypothesis generation through an AD application.

A key scientific contribution of FeatureMAP is the integration of uncertain metabolite annotations with network-guided discovery of metabolic dysregulation. Building on mummichog, PIUMet, and our previous TidyMass2 framework, FeatureMAP uses metabolic network connectivity to refine ambiguous feature-metabolite assignments and prioritize candidates supported by coherent biochemical relationships^3,4,8^. Importantly, network-based prioritization provides complementary evidence rather than definitive metabolite identification. Beyond annotation refinement, FeatureMAP offers an alternative to conventional pathway enrichment, which relies on predefined pathway boundaries and often treats metabolite changes analogously to gene expression^5,19^. However, metabolite abundances are governed by interconnected processes of synthesis, consumption, and transport and do not necessarily reflect pathway activity^5,6^. By integrating continuously weighted feature-level evidence with metabolic reaction networks, FeatureMAP identifies data-driven modules that can span conventional pathway boundaries and incorporate unobserved connecting metabolites. These modules provide a complementary representation of metabolic perturbations that is not restricted to confidently identified metabolites or predefined pathways.

The biological interpretation of network-derived modules presents a further challenge, as individual modules may contain metabolites associated with multiple biochemical processes or misleading network connections. To address this, FeatureMAP combines structured biochemical evidence with LLM-based functional interpretation. The fast mode provides standardized biochemical annotations, whereas the advanced mode uses multi-agent hypothesis generation, critique, revision, and arbitration to evaluate competing interpretations. Our benchmarks showed that fast mode generally recovered the underlying biochemical themes despite substantial metabolite-level noise, although detailed functional summaries were more sensitive to competing signals. Advanced mode improved a subset of difficult interpretations, highlighting the potential of structured critique to reduce misleading functional assignments. Nevertheless, LLM-generated interpretations remain dependent on the underlying biochemical evidence and should be regarded as hypotheses rather than experimentally established mechanisms^20^.

FeatureMAP-derived modules provide a common biochemical representation for comparing studies with different analytical coverage. PhenoMAP integrates these modules with ontology-normalized phenotypes and associated metabolites in a unified knowledge graph. Greater similarity between independent studies of the same phenotype, together with concordance with curated disease-pathway associations, supports the reproducibility and biological relevance of module-based comparisons. The stable phenotype communities further illustrate how this representation can connect diseases, exposures, and interventions through related metabolic processes. For example, arachidonic-acid-related modules linked smoking, asthma, cellular senescence and methylprednisolone-associated responses, consistent with established roles of eicosanoids in smoking-associated airway inflammation, senescence and glucocorticoid responses^21–23^. Similarly, the lipid-related connections among renal and hepatic diseases agree with evidence of disrupted fatty-acid metabolism in kidney fibrosis and steatotic liver disease–associated hepatocellular carcinoma^24,25^. These examples support using PhenoMAP to identify biochemical links among heterogeneous phenotypes and prioritize processes for cross-context investigation. Although such links do not establish identical molecular changes, common causal mechanisms or therapeutic reversal, the retained modules, metabolites, and study contexts provide the evidence needed to formulate and test these possibilities.

The AD application illustrates how cross-system module convergence can connect established lipid pathology with specific hypotheses for experimental investigation. The membrane-lipid modules identified across CSF and AD-derived iPSCs were consistent with previous evidence implicating ceramide and sphingolipid metabolism in AD^26,27^. The convergent galactose-related modules further highlighted ceramide-derived glycosphingolipids, including Gb2 in both systems, sulfatide in CSF, and Gb3 in iPSCs. This candidate pattern focuses attention on glycosphingolipid headgroup metabolism and lysosomal turnover: Gb2 and Gb3 share A4GALT-mediated galactosylation and GLA-dependent degradation, providing concrete enzymatic entry points for follow-up studies^28^. These relationships nominate altered synthesis or clearance of galactosylated glycosphingolipids as a potential component of AD-associated membrane remodeling. A possible functional consequence is altered glial lipid handling and immune regulation. Experimental depletion of myelin sulfatide induces disease-associated microglial and astrocytic responses and cognitive impairment, establishing that disruption of a specific glycosphingolipid can provoke AD-like neuroinflammation^29^. The additional arachidonic-acid-related modules identified in both datasets provide a complementary lipid-signaling context, supported by experimental links between leukotriene metabolism and microglia in AD models^30^. Although these observations do not establish a microglial origin or causal role for our candidates, they motivate testing whether glycosphingolipid remodeling affects lipid clearance and inflammatory responses in relevant neural-glial systems.

Several limitations should be considered. First, FeatureMAP depends on metabolic reaction networks that remain incomplete, particularly for tissue-specific, compartmentalized, and poorly characterized biochemical processes, potentially limiting the recovery of certain metabolic modules. Second, network-refined metabolite assignments remain putative and require orthogonal structural validation. Third, LLM-based interpretation may introduce unsupported hypotheses or model-dependent biases, emphasizing the importance of evidence traceability and independent biological evaluation. In addition, PhenoMAP comparisons remain susceptible to differences in biospecimens, analytical platforms, and study designs, while the AD findings require validation in larger independent cohorts and experimental models. Furthermore, the iPSC analysis included only two independent donor-derived cell lines per group, with three culture replicates per line, potentially introducing pseudo-replication and limiting the statistical robustness and generalizability of the findings.

Future development should focus on expanding metabolic network coverage, incorporating tissue- and context-specific biochemical information, integrating complementary omics data, and strengthening experimental validation of predicted metabolic relationships. Continued expansion of PhenoMAP may further enable systematic investigation of metabolic dysregulation across human phenotypes. Together, FeatureMAP and PhenoMAP provide an integrated framework for transforming incompletely annotated LC-MS data into interpretable metabolic modules and using these representations to generate testable biological hypotheses.

## Methods

### Databases in FeatureMAP

*Metabolite reference database*. The metabolite reference database used for MS1-based annotation was constructed by integrating compound libraries from the HMDB and KEGG human databases^31,32^. Initially, 217,920 HMDB and 24,002 KEGG compounds were retrieved. Redundant KEGG entries were removed based on cross-referenced HMDB identifiers, retaining 9,969 unique KEGG compounds. The resulting database comprised 227,889 compounds, with reference *m/z* values, molecular formulas, and database identifiers where available^33–35^.

#### Human metabolic reaction network

The human metabolic reaction network (MRN) was constructed by integrating curated reaction information from nine public resources: KEGG, HMDB, Recon3D, ModelSEED, MetaNetX, BiGG, Rhea, Reactome, and HumanCyc (Supplementary Table 4)^31,32,36–42^. For each database, the full reaction tables were downloaded from the original repositories and parsed to obtain metabolite-metabolite reaction pairs, treating each biochemical reaction as a set of substrate-product pairs^3,43^. Metabolite identities were harmonized across resources using standardized InChI identifiers, and duplicate reaction pairs referring to identical reactant-product combinations were removed during the merging process. The resulting MRN comprises 9,730 metabolite nodes and 31,666 reaction edges, representing, to our knowledge, one of the most extensive human metabolic reaction networks reported to date, and was stored as a graph object in R for all subsequent analyses.

### Raw feature annotation via evidence-integrated iterative scoring

Raw LC-MS features were first annotated using an evidence-integrated MS1 workflow adapted from our previous TidyMass2 framework^3^. The input feature table contains, for each feature, *m/z*, retention time (RT), ionization mode, mean intensity, effect size, and P value. Features were matched by *m/z* using polarity- and column-specific adduct rules within a user-defined mass tolerance, and all matched metabolites were retained as candidate annotations. Candidate annotations of the same metabolite were grouped into metabolite feature clusters (MFCs), each comprising RT-coherent features that support the metabolite as the monoisotopic ion, isotope peaks, or alternative adducts. Features of the same metabolite eluting at distinct retention times formed separate MFCs.

Each MFC was assigned an evidence score integrating adduct consistency, isotopic evidence, and, when available, MS/MS annotations (Supplementary Methods). Redundant annotations were reduced through iterative pruning and rescoring at the metabolite and feature levels. Lower-scoring candidates were removed only when higher-scoring competing annotations were present, and the procedure continued until no further reduction in redundancy was achieved (Supplementary Methods). Each retained feature-metabolite assignment inherited the evidence score of its corresponding MFC, which was divided by the highest evidence score among all retained assignments in the same dataset, yielding a normalized annotation score 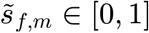 for feature *f* and metabolite *m*. The resulting candidate annotations, referred to as MS1-based annotations, served as input to network refinement.

### Inference of a feature-enriched metabolic reaction subnetwork via PCSF

Even after redundancy reduction, the MS1-based annotations may still contain multiple plausible candidates for some features^3^. Previous evidence has shown that genuine metabolic perturbations tend to form topologically coherent clusters within the reaction network, whereas spurious assignments remain isolated or weakly connected^3,4,8^. The PCSF framework was therefore applied to prioritize biologically consistent annotations based on network connectivity^8^. Unlike approaches that rely on predefined significance thresholds, all detected features were included by default and weighted using a continuous importance score (defined below). PCSF identifies a subnetwork that maximizes the total node prize while minimizing edge costs^44^. For PCSF input, each feature in the MS1-based annotations was represented as a node and connected to its candidate metabolites by feature-metabolite edges. These edges, together with metabolite-metabolite reaction edges from the MRN, defined the feature-metabolite graph for PCSF optimization.

#### Definition of node prize

Node prizes were specified separately for features and metabolites. For each feature *f*, an importance score *I*_*f*_ was defined as

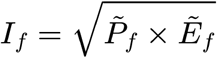

Where *P*_*f*_ = −long_10_(*pf*) and E*f* =|*S*_*f*_ |. Here, *p* _*f*_ denotes the P value from the association test for feature *f*, and *S*_*f*_ represents an effect size measure. For categorical outcomes (*e*.*g*., case-control comparisons), *S*_*f*_ is given by log_2_, whereas for continuous outcomes (*e*.*g*., quantitative traits), *S*_*f*_ corresponds to either the regression coefficient or the correlation coefficient. To place statistical significance and effect size on a comparable scale, both *P*_*f*_ and *E*_*f*_ were rescaled to the [0, 1] interval using min-max normalization across all features, yielding 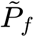 and 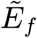, respectively. Only features that are both significant and of substantial magnitude therefore receive high importance scores. The prize of each feature node was set to its importance score, *π* _*f*_ = *I*_*f*_.

For metabolite nodes, a degree-based penalty was imposed to discourage the selection of highly connected hubs that could artificially link unrelated regions of the network. Specifically, the prize assigned to a metabolite node *m* was defined as

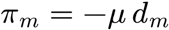

where *d*_*m*_ denotes the degree of metabolite *m* in the feature-metabolite graph, and the scaling parameter *μ* keeps the degree-based penalty small relative to feature prizes (Supplementary Methods).

#### Definition of edge cost

Edge costs were defined to reflect annotation confidence and network structure. For feature-metabolite edges, the cost was set to

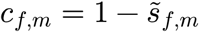

Where 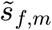 is the annotation score of the corresponding candidate annotation. Under this formulation, a lower annotation score results in a higher edge cost, making such connections less likely to be retained by PCSF. For metabolite-metabolite edges, which represent curated chemical reactions, a constant cost equal to the maximum feature-metabolite edge cost was assigned, following previous studies (Supplementary Methods)^8^.

#### PCSF optimization

With node prizes and edge costs specified as above, the PCSF algorithm, as implemented in the R package PCSF, was applied to the feature-metabolite graph^10^. PCSF identifies an optimal forest *F* with node set *V*_*F*_ and edge set *E*_*F*_ by minimizing

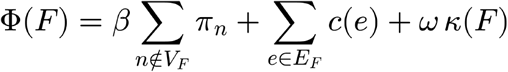

where *π*_*n*_ is the prize of node *n*,^*c*(*e*)^, is the cost of edge *e*, and *k*(*F*) is the number of trees in *F*. The three terms penalize, respectively, the exclusion of high-prize nodes, the inclusion of costly or weakly supported edges, and excessive fragmentation of the forest, with β and ω controlling their relative weights (Supplementary Methods). In addition to pruning unreliable edges, PCSF incorporates metabolites without a connected feature, termed hidden metabolites, when they provide the most efficient routes between high-prize regions that would otherwise remain isolated.

#### Construction of a robustness-weighted PCSF subnetwork

Given that a single PCSF solution may be sensitive to small variations in edge costs and represents only one of many plausible subnetworks, the robustness of nodes and edges was assessed by introducing random multiplicative perturbations to the edge costs and repeating the optimization. For each replicate (*r*=1,…,*R*(*R*=100)by default), edge costs were perturbed as

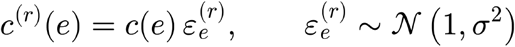

Where 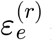 is drawn independently for each edge and replicate, and *σ* =0.2 by default. PCSF was then applied to the graph with costs *c*^(*r*)^ (*e*), yielding a forest *F*^(*r*)^ with node set 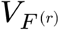 and edge set 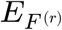.

Robustness was quantified by the node and edge frequencies

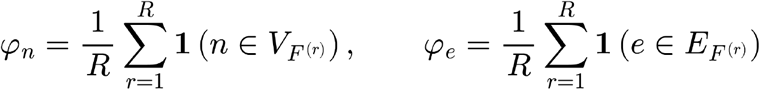

which give the proportion of replicates in which node or edge is included in the PCSF solution. The union of all forests 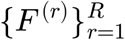, together with the associated node and edge frequencies, defines the PCSF subnetwork used in downstream analyses. Candidate annotations whose feature-metabolite edges did not appear in any of the forests were removed, and the retained candidate annotations are referred to as network-refined annotations. Each network-refined annotation was assigned a network-weighted annotation score combining its annotation score with the frequency of its feature-metabolite edge,

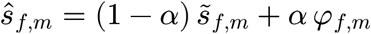

where *φ*_*f,m*_ is the edge frequency of the feature-metabolite edge and the weight is adapted to the variability of edge frequencies in each dataset (Supplementary Methods).

### Identification of functional modules via community detection

Although PCSF retains topologically coherent signals, the PCSF subnetwork may still span multiple biological processes as well as residual noise. Because dysregulated metabolic processes are expected to form densely connected regions of the reaction network, whereas unrelated metabolites remain sparsely connected, community detection was applied to partition the subnetwork into modules defined by reaction connectivity rather than predefined pathways^3,45^.

#### Graph-based community detection

Metabolites retained in the PCSF subnetwork were mapped back onto the MRN, and the subgraph induced by these metabolites was used for community detection (Supplementary Methods). Each reaction edge between metabolites and was weighted by its edge frequency, normalized by the degrees of its endpoints in the MRN to attenuate hub-driven aggregation:

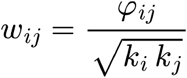

where *k*_*i*_ is the degree of metabolite *i* in the MRN. Communities were identified on this weighted subgraph using the Louvain algorithm by default^46^. The leading eigenvector, fast greedy, and Walktrap algorithms are provided as alternatives^47–49^.

#### Calculation of module score

Candidate modules were scored using a modularity-based activity measure adapted from mummichog and extended to capture the magnitude of dysregulation. For a module *G* with *N*_*I,G*_ metabolites, of which are annotated metabolites, that is, metabolites connected to at least one feature in the PCSF subnetwork, the baseline activity score was defined as

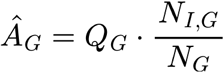

where is *Q*_*G*_ the adjusted Newman-Girvan modularity of computed on the MRN^50^. Each annotated metabolite was assigned a metabolite-level importance *I*_*m*_, computed as the mean importance score of its features weighted by their network-weighted annotation scores (Supplementary Methods). The module score was then defined as

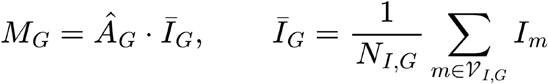

where *V* _*I,G*_ is the set of annotated metabolites in *G*, so that high-scoring modules are topologically coherent, enriched for annotated metabolites and composed of strongly dysregulated metabolites.

#### Evaluation of module significance

The significance of each candidate module was assessed against a degree-matched permutation null, in which community detection and module scoring were repeated on randomized MRN subnetworks (Supplementary Methods). Candidate modules with FDR < 0.05 were designated as dysregulated modules.

### Dual-mode agentic LLM framework for functional module interpretation

Each dysregulated module was interpreted by a framework that combines a structured evidence profile with LLM reasoning. The framework comprises two modes: a fast mode for standardized annotation and an advanced mode that emulates peer review, in which candidate interpretations are proposed, challenged, revised, and adjudicated.

#### Module evidence profile

Metabolites in each module were mapped to HMDB and KEGG records to obtain standardized names, identifiers, and chemical classes. Metabolites were ranked by a priority score combining metabolite-level importance and node frequency, and prioritized metabolites were selected adaptively from the distribution of this score (Supplementary Methods). Prioritized metabolites served as the primary evidence for interpretation, whereas all module metabolites were summarized by chemical class to retain compositional context. The profile further included signal concentration, a Herfindahl-Hirschman-type index indicating whether the dysregulation signal is concentrated in a few metabolites or distributed across many; pathway coverage derived from KEGG and HMDB pathway records; and an origin profile giving the proportions of endogenous, exogenous, and unknown-origin metabolites (Supplementary Methods).

#### Fast mode

Fast mode separated module-intrinsic interpretation from study-context evaluation using two sequential LLM calls. The first call received only the module evidence profile, with species, biospecimen, and phenotype deliberately excluded to prevent context-driven overinterpretation, and returned four fields: origin type (Host-Endogenous, Microbiome-Derived, Diet-Derived, Xenobiotic/Exogenous, or Host-Microbiome), functional title, functional summary, and anchor metabolites, defined as the prioritized metabolites most directly supporting the interpretation. The second call evaluated this interpretation in the study context using the origin profile, species, biospecimen, phenotype, and, optionally, PubMed abstracts retrieved with the functional title and anchor metabolites (Supplementary Methods). It returned a phenotype relationship, a confidence label (high, medium, or low) describing the strength of evidence for the interpretation, and a supporting rationale. Both calls returned structured JSON outputs.

#### Advanced mode

Advanced mode formulated interpretation as a multi-agent peer review with role-specific access to information (Supplementary Methods)^51^. A Proposer generated one to three candidate hypotheses from the module evidence profile alone, each comprising an origin type, functional title, functional summary, anchor metabolites, and, optionally, tension metabolites that are difficult to reconcile with the proposed mechanism. A Skeptic critiqued each hypothesis using the module evidence, origin profile, species, and biospecimen, but without access to phenotype or literature, and assigned each critique to a predefined category, such as *weak metabolite support, generic hub bias, isolated outlier bias*, or *biochemical conflict* (Supplementary Methods and Supplementary Table 5). Each critique was graded as *major* (if it undermined the central mechanism), *minor* (if it warranted acknowledgment only), or no challenge (if no substantive concern was identified). A Reviser then kept, modified, or dropped each hypothesis, and a second Skeptic-Reviser round was triggered when substantive revisions or unresolved major critiques remained. Literature evidence was then retrieved and summarized for each surviving hypothesis, and these summaries, together with the debate history and study context, were supplied only to an Arbiter, which accepted or rejected each hypothesis and assigned a confidence label with rationale.

#### Cross-module synthesis

In advanced mode, accepted hypotheses across modules can optionally be integrated by a synthesis agent into thematic clusters and an overall summary. Biology links between hypotheses were inferred by the agent from their functional titles and summaries, whereas biochemical links were computed independently of the LLM, requiring a direct MRN reaction edge between prioritized metabolites and a shared pathway. Each pair of hypotheses was classified as biochemical, biology, both, or none (Supplementary Methods).

#### Interpretation outputs

Both modes returned structured records comprising database-derived context (chemical-class composition, pathway coverage, and origin profile), the interpretation (origin type, functional title, functional summary, and anchor metabolites), the phenotype relationship, confidence label and rationale, and global direction, a statistical descriptor of the overall up- or downregulation of the module (Supplementary Methods). Advanced mode additionally retained the complete evidence trail, including intermediate agent outputs, critiques, revisions, and verdicts, together with the synthesis network when cross-module synthesis was enabled.

### Evaluation of network-guided MS1 annotation refinement

#### Real-data cohort selection and processing

Ten case-control untargeted LC-MS metabolomics datasets were selected from MetaboLights, spanning diverse biospecimen types (cell culture, circulating cells, plasma/serum, solid tissue and feces) and biological contrasts^16^. All datasets were processed using the TidyMass pipeline, including feature detection, alignment, differential analysis, and MS/MS spectral matching against the built-in public spectral libraries^33^. For each feature, the top-scoring MS/MS match was retained as the reference annotation, and a candidate annotation was considered correct when it matched this reference. MS/MS spectra were excluded from the FeatureMAP analysis to ensure an independent and unbiased evaluation of annotation performance.

#### Annotation performance metrics

Annotation performance was evaluated on the top-1 to top-3 ranked candidates of each feature using three metrics (Supplementary Methods). Tie-adjusted annotation accuracy measures whether the correct annotation is ranked among the top candidates, discounted by the number of candidates tied with it. The F1 score measures whether correct annotations are recovered without excessive incorrect candidates. Annotation redundancy, the mean number of candidate annotations per feature, measures how specific the annotation output is.

#### Benchmarking against mWISE and xMSannotator

mWISE and xMSannotator were selected as comparators because, like FeatureMAP, both combine *m/z*, retention time, and intensity information to annotate MS1 features (Supplementary Table 6)^11,12^. All three tools were applied to the same ten metabolomics datasets, and evaluation was restricted to features with an MS/MS-supported reference annotation that were annotated by all three tools, providing a common test set.

### Spike-in simulation benchmark for module identification

#### Semi-synthetic benchmark design

A spike-in benchmark was constructed using 40 control participants from a gestational diabetes cohort (MTBLS12326), who were randomly split into pseudo-case and pseudo-control groups (n = 20 each)^13^. Arginine biosynthesis (KEGG ID: hsa00220) was selected as the target pathway, and its seven metabolites with MS/MS-supported annotations, corresponding to 11 features, were designated as spike-in targets (Supplementary Data 2). In each of 100 simulation runs, the intensities of these features in the pseudo-case group were scaled by feature-specific fold changes drawn uniformly from 2 to 10 (Supplementary Methods). Pathway metabolites whose features were scaled were defined as spiked metabolites, and the remaining pathway metabolites in the MRN were defined as unspiked metabolites. A run was considered successful if exactly one dysregulated module was identified and it contained at least five spiked metabolites. To evaluate whether FeatureMAP can resolve concurrent perturbations, the benchmark was extended to a two-pathway setting in which arginine biosynthesis and biosynthesis of unsaturated fatty acids (KEGG ID: hsa01040) were spiked simultaneously, comprising 25 spiked metabolites (7 and 18, respectively) and 43 LC-MS features, across 100 runs with the same background and design (Supplementary Data 2 and Supplementary Methods).

#### Benchmarking against mummichog

FeatureMAP was compared with *mummichog* using the same 100 simulated feature tables (Supplementary Methods and Supplementary Table 6). For both tools, the module with the lowest P value in each run was defined as the primary module. Recovery was quantified as the number of spiked and unspiked metabolites in the primary module and compared between tools using paired Wilcoxon signed-rank tests. Success rate was defined as the proportion of runs in which the primary module contained at least k spiked metabolites (k = 1, …, 7) and was the only significant module in that run (FDR < 0.05 for FeatureMAP and nominal P < 0.05 for mummichog, following the default output of each tool).

#### Random metabolite negative control

To assess specificity, seven MRN-mapped metabolites with MS/MS-supported annotations were randomly selected in each of 100 runs, and their features were spiked as described above. The proportion of runs yielding at least one dysregulated module was recorded, and the FDR and module scores of these modules were compared with those from the pathway spike-in using two-sided Wilcoxon rank-sum tests.

### Evaluation of dual-mode LLM functional annotation

#### Construction of semi-synthetic metabolic modules

To benchmark functional annotation against known biochemical pathways, semi-synthetic metabolic modules were constructed using curated human pathways from PathBank (Supplementary Data 3)^14^. Pathways were included if more than 50% of their HMDB-annotated metabolites could be mapped to the MRN, and the mapped metabolites served as the ground-truth signal set for each pathway. Neighbor metabolites were then sampled within 1-5 reaction steps of the pathway metabolites, and noise metabolites within the same range of the neighbor metabolites, each equal in number to the pathway metabolites and without overlap among the three sets. For each of the 104 eligible pathways, one module was generated in an easy and in a hard setting, yielding 208 modules in total. Pathway metabolites were assigned high metabolite-level importance and node frequency values, whereas neighbor metabolites received intermediate values. Noise metabolites were assigned low values in the easy setting but values comparable to those of pathway metabolites in the hard setting, creating competing signals that challenged the identification of the underlying pathway (Supplementary Methods).

#### Evaluation of fast-mode annotation

All modules were annotated using fast mode with gpt-4.1-mini, without phenotype or literature input. Generated titles were evaluated against the reference pathway names in three ways. First, title similarity was quantified as the cosine similarity of OpenAI, SapBERT, and MedCPT embeddings and compared with that of randomly shuffled, non-matching title pairs^52,53^. Second, titles were classified as exact matches, partial word overlaps, or no word overlap with the reference name. Third, for titles with partial word overlap, ROUGE-L recall was used to quantify the fraction of the reference name recovered by the generated title (Supplementary Methods)^15^. Generated functional summaries were evaluated against the reference pathway descriptions using OpenAI and MedCPT embedding similarity and an LLM-as-judge framework that scored topic alignment, entity support, mechanistic consistency, and absence of unsupported claims (Supplementary Table 2). A reverse judge additionally scored the severity of topic deviation, unsupported entities, mechanistic inconsistency, and hallucinated claims (Supplementary Methods and Supplementary Table 3).

#### Evaluation of advanced-mode rescue

Modules with weak fast-mode title recovery, defined as those with no word overlap or with partial word overlap and a ROUGE-L recall below the median of partial-overlap modules in the same difficulty setting, were re-annotated using advanced mode. A module was considered rescued when a title with no word overlap achieved partial or exact overlap, or when a partial-overlap title achieved a higher ROUGE-L recall. For rescued modules, LLM-as-judge scores of fast- and advanced-mode summaries were compared using two-sided Wilcoxon signed-rank tests.

### Construction of the PhenoMAP knowledge graph

#### Public study curation

PhenoMAP incorporates public human untargeted LC-MS metabolomics studies from Metabolomics Workbench and MetaboLights^16,17^. Eligible studies were required to provide MS1-level raw data and sample metadata sufficient to define group comparisons. MS/MS spectra were used when available but were not required. Studies with two groups were analyzed as case-control comparisons, and studies with multiple groups or factors were decomposed into pairwise comparisons of each group against a reference group. Raw data were processed using the TidyMass pipeline, differential analysis was performed for each comparison, and FeatureMAP modules were annotated using fast mode with Qwen3.5-122B-A10B. Studies yielding at least one dysregulated module were retained, resulting in 501 studies.

#### Phenotype normalization and ontology mapping

For each study, the repository study description was used to extract the principal phenotype with GPT-4.1, which assigned it to one of nine categories: disease, symptom, intervention, chemical, exposure, trait, cell type/line, genetic perturbation, or biological process. Each phenotype was then mapped to ontology terms within category-specific ontology sources (Supplementary Methods). Ambiguous, missing, or biologically inconsistent mappings were manually reviewed before graph construction.

#### Graph entities and relationships

PhenoMAP was organized as a heterogeneous knowledge graph with four node types: studies, which retain repository provenance and experimental context; phenotypes, which represent ontology-normalized biological or clinical variables; modules, which represent FeatureMAP-derived dysregulated modules; and metabolites, which represent metabolites in the MRN. Studies were connected to phenotypes by “investigated” relationships and modules to studies by “reported in” relationships. Metabolites were connected to modules by “member of” relationships labeled as core (anchor metabolites), supporting (other prioritized metabolites), or contextual (remaining module metabolites), and to one another by “connected with” relationships derived from MRN reactions. To enable cross-study comparison, module pairs were scored by combining the similarity of their functional annotations with the overlap of their member metabolites, chemical classes, and pathway coverage, and pairs with a similarity score of at least 0.7 were connected by “similar to” relationships (Supplementary Methods).

#### Graph-enabled analysis

PhenoMAP provides four analysis functions built on the knowledge graph (https://phenomap.jaspershenlab.com/). (1) *Entity Search* retrieves studies, phenotypes, modules, and metabolites by name, keyword, external database identifier, or stable PhenoMAP identifier, and links each entity to a node-detail page reporting its properties and neighboring records. (2) *Path Query* identifies connections between two entities by returning all study, phenotype, and module nodes on paths up to a user-defined maximum length. (3) *Module Enrichment* was designed for metabolite lists obtained outside the FeatureMAP workflow, such as differential metabolites from annotated untargeted data or from targeted assays, and tests whether such a list is over-represented in PhenoMAP modules without requiring feature-level reanalysis. Each module was represented by its core and supporting metabolites, and the background was defined as the union of these metabolites across all modules. Enrichment was assessed using a one-sided hypergeometric test with Benjamini-Hochberg correction, and modules with FDR < 0.05 and at least two overlapping metabolites were reported as enriched by default (Supplementary Methods). (4) *PhenoMAP Chat* provides a natural-language interface to these functions: each question is assigned to a query type, the corresponding PhenoMAP function is called, and the response is generated from the returned records, with every cited study, phenotype, module, and metabolite linked to its PhenoMAP identifier (Supplementary Methods). Chat is available through user-provided OpenAI or SiliconFlow API credentials. A community submission workflow accepts user-contributed studies, which are reviewed and processed through the same pipeline before inclusion in future releases (Supplementary Methods).

#### PhenoMAP platform implementation

The knowledge graph was stored in Neo4j, and entity search, node-detail retrieval, path query, and graph summary functions were served through a Go backend API. Module enrichment was provided by an R Plumber service, and PhenoMAP Chat was implemented as a Python FastAPI service connected to the same graph and enrichment APIs. The user interface was built with Vue 3 and Vite. User authentication, downloads, and community submissions were handled by the backend, with submitted files stored separately from the public graph.

### Cross-study analyses in PhenoMAP

#### Phenotype similarity

Cross-study analyses used high-confidence Host-Endogenous modules, which reflect intrinsic host metabolism and are comparable across studies. Similarity between two phenotypes or studies, each represented by a set of modules, was quantified as the BMA: each module was matched to its most similar module in the other set, and the best-match similarities were averaged in both directions (Supplementary Methods)^18^.

#### Validation of phenotype similarity

Two analyses tested whether module-based similarity captures reproducible and biologically meaningful relationships. First, to assess cross-study reproducibility, all modules of one phenotype from one study were treated as a study-phenotype unit. For phenotypes investigated in at least two studies, BMA similarity was calculated between units from different studies, and similarities between units of the same phenotype were compared with those between units of different phenotypes.

Second, to assess biological validity, disease phenotypes were mapped to the CTD, and their curated associations with KEGG pathways, which focus on metabolism, were retrieved^54^. For each pair of diseases, the number of KEGG pathways associated with both diseases was counted, and pairs sharing more pathways than the average across all pairs were defined as sharing a metabolic mechanism. BMA similarity was then compared between disease pairs with and without a shared metabolic mechanism. Both comparisons used one-sided Wilcoxon rank-sum tests.

#### Phenotype communities

To identify groups of phenotypes with related metabolic dysregulation, a phenotype network was constructed by connecting the most similar phenotype pairs, and communities were identified using consensus Louvain clustering. Community robustness was assessed by bootstrap resampling of phenotypes and summarized using the mean bootstrap Jaccard similarity (Supplementary Methods)^55,56^. To assess whether the connections in these communities reflect known biology, each pair of connected phenotypes was checked against the literature. A connection was considered directly supported when published studies reported an association between the two phenotypes, and indirectly supported when no such study existed but both phenotypes were independently linked to the same specific biological mechanism (Supplementary Methods).

### Clinical cohort and CSF collection

The clinical cohort comprised 55 patients with AD and 21 non-AD clinical controls recruited at the First Affiliated Hospital, Zhejiang University School of Medicine, Hangzhou, China. Patients with AD were consecutively recruited from the Memory Clinic of the Department of Neurology. AD was diagnosed according to the 2011 National Institute on Aging-Alzheimer’s Association (NIA-AA) core clinical criteria for all-cause dementia and mild cognitive impairment, based on clinical history, neurological examination, neuropsychological assessments, brain imaging, and available AD-related biomarker evidence^57,58^. Neuropsychological assessments included the Mini-Mental State Examination, Montreal Cognitive Assessment, Clock Drawing Test, Auditory Verbal Learning Test, and Clinical Dementia Rating. The non-AD clinical controls comprised patients with other neurological or medical conditions. For AD cases, exclusion criteria included alternative causes of cognitive impairment, severe malnutrition or infection, recent drug or alcohol abuse, major psychiatric disorders, severe systemic diseases, and sensory or motor impairments interfering with cognitive assessment. The study was approved by the Ethics Committee of the First Affiliated Hospital, Zhejiang University School of Medicine (approval number: 2025-898). Written informed consent was obtained from all participants or their legally authorized representatives before sample collection. CSF samples were collected from all participants by lumbar puncture into polypropylene tubes. Following collection, CSF samples were centrifuged and stored at −80°C until untargeted metabolomic analysis.

To minimize potential confounding by age and sex, 21 patients with AD were selected from the clinical cohort and individually matched to the 21 non-AD controls at a 1:1 ratio. Matching required identical sex and was optimized to minimize the overall absolute age differences between matched pairs, without replacement. This process yielded 21 age- and sex-matched AD-control pairs for subsequent analyses.

### iPSC culture and sample collection

Fibroblasts from two individuals with sporadic AD (sAD; AG06869 and AG08243) and two healthy controls (AG04148 and AG08525) were obtained from the Coriell Cell Repository and reprogrammed into iPSCs as previously described^59^. iPSCs were maintained on Matrigel-coated dishes (Corning, 354237) in complete mTeSR medium (STEMCELL Technologies), with medium replacement every other day. Cells were passaged using dispase (STEMCELL Technologies) at approximately 50% confluence. Three independently cultured replicates were prepared from each iPSC line, yielding six samples per group from two donor-derived lines. Each replicate was harvested as a separate cell pellet and processed independently using the same protocol.

### Untargeted LC-MS metabolomic profiling and data processing

CSF aliquots (100 μL) were extracted with 400 μL of pre-cooled methanol–acetonitrile (1:1, v/v) containing internal standards. iPSC pellets containing 1 × 10^6^ cells per sample were extracted with 200 μL of methanol-acetonitrile-water (2:2:1, v/v/v). Samples were vortexed, with iPSC samples additionally subjected to three liquid-nitrogen freeze–thaw cycles. Both sample types were sonicated in an ice-water bath for 10 min and incubated for 1 h at -20 °C. Extracts were then centrifuged at 13,800 × g for 15 min at 4 °C, and supernatants were transferred to autosampler vials. Pooled quality-control (QC) samples were prepared by combining 10 μL aliquots from each sample extract.

Extracts were analyzed using a Vanquish ultra-high-performance liquid chromatography system coupled to an Orbitrap Exploris 120 mass spectrometer (Thermo Fisher Scientific). Chromatographic separation was performed using BEH Amide (Waters) and Kinetex C18 (Phenomenex) columns for HILIC and RP analysis, respectively, with data acquired in positive and negative electrospray ionization modes. Pooled QC samples and blanks were included to monitor analytical stability and carryover. Raw data were converted to mzXML format using ProteoWizard and processed using an in-house XCMS-based R pipeline for peak detection, alignment, and integration^33,60,61^. Metabolites were annotated by matching MS/MS spectra against BiotreeDB v4.0, with annotation confidence classified according to the Metabolomics Standards Initiative (MSI) criteria^62^.

### Cross-system analysis of AD-associated metabolic modules

CSF and iPSC datasets were analyzed independently using FeatureMAP, with HILIC and RP data processed separately. Feature-level differences were assessed using two-sided paired Wilcoxon signed-rank tests for matched CSF samples and Wilcoxon rank-sum tests for iPSC samples. Candidate modules with nominal P < 0.05 were interpreted using advanced mode, and accepted hypotheses classified as Host-Endogenous with high interpretation confidence were retained for cross-system comparison. CSF and iPSC annotations were compared using the module-module similarity defined above. Pairs with similarity scores ≥ 0.70 were retained, and the corresponding functional annotations were manually grouped into metabolic themes based on their titles and summaries.

To assess the extent of AD-related literature associated with the shared metabolic themes, we performed module-title-based searches in PubMed. For each module, PubMed was searched using its functional title together with AD-related terms. Retrieved records were deduplicated by PMID within each module. Counts were then summed across modules within each metabolic theme and divided by the number of modules to obtain the mean number of retrieved publications per module. Results were summarized as mean ± standard deviation.

For the membrane-lipid-remodeling and galactose-metabolism examples, anchor candidates from the corresponding CSF and iPSC module annotations were manually organized into biochemical maps using KEGG reaction and pathway definitions, with connections checked against the MRN where available. The maps distinguished shared and system-specific candidate support from connecting intermediates.

For comparison with conventional metabolite-based analysis, metabolites with vendor-provided Level 1/2 annotations were evaluated using two-sided paired Wilcoxon signed-rank tests for matched CSF samples and two-sided Wilcoxon rank-sum tests for iPSC samples. In each dataset, metabolites with nominal P < 0.05 were subjected to KEGG pathway enrichment analysis. Additionally, Level 1 annotations in both datasets were used as references to evaluate top 1-3 tie-adjusted annotation accuracy, following the procedure described above. Annotation redundancy was calculated at each annotation stage as the mean number of candidate annotations per feature.

## Code availability

All data processing and statistical analyses were performed using R version 4.5.2, Python 3.9.6 and associated packages. The source code for FeatureMAP and FeatureMAPShiny is available on GitHub at https://github.com/tidymass/featuremap and https://github.com/tidymass/featuremap_shiny, respectively. The source code for PhenoMAP is at https://github.com/tidymass/phenomap. Scripts used for data processing, statistical analysis, and figure generation are available at https://github.com/tidymass/featuremap_manuscript. Detailed tutorials are provided at https://www.tidymass.org/featuremap_tutorial. The interactive FeatureMAP and PhenoMAP web applications are accessible at https://featuremap.jaspershenlab.com and https://phenomap.jaspershenlab.com, respectively.

## Data availability

Public metabolomics datasets reanalyzed in this study were obtained from MetaboLights (https://www.ebi.ac.uk/metabolights/) and Metabolomics Workbench (https://www.metabolomicsworkbench.org/), with study accession numbers and associated metadata provided in the Supplementary Information. The CSF and iPSC metabolomics datasets generated in this study are available upon reasonable request to X.S., G.P., and L.Z. Processed results supporting the findings of this study are provided in the Supplementary Data.

## Acknowledgments

This work was supported by start-up funding associated with the Nanyang Assistant Professorship awarded to Xiaotao Shen by the Lee Kong Chian School of Medicine and the School of Chemistry, Chemical Engineering and Biotechnology at Nanyang Technological University, Singapore. Additional funding was provided by a Ministry of Education (MOE) Tier 1 grant awarded to Xiaotao Shen (Grant No. 025402-00001). Chuchu Wang was supported by start-up funding through the Nanyang Assistant Professorship from the Lee Kong Chian School of Medicine, Nanyang Technological University, Singapore. The authors acknowledge the use of ChatGPT (OpenAI) for language editing and grammatical refinement during manuscript preparation.

## Author Contributions

X.S. conceived the project and supervised the study. Y.W. and Y.L. modified the MS1-based metabolite annotation workflow. Y.W. and X.S. developed the FeatureMAP algorithm and software. Y.W. and F.Z. developed PhenoMAP. G.P. coordinated the preparation of the AD CSF samples and obtained IRB approval. L.Q. and L.Z. prepared the iPSC samples. Y.W. performed all metabolomics data processing, analysis, and case-study investigations. Y.W. and X.S. prepared the figures. X.S. and Y.W. wrote the manuscript. J.K., X.Y., C.L., C.J., Q.S., and C.W. contributed to manuscript revision. All authors approved the final version.

## Competing Interests

The authors declare no conflict of interest.

## Additional information

Correspondence and requests for materials should be addressed to X.S.

## Notes

### Competing Interest Statement

The authors have declared no competing interest.

