## Supplementary Information for "FeatureMAP enables network-guided interpretation of untargeted metabolomics"

### Table of Contents

#### Supplementary Figures


1. **Supplementary Fig. 1.** Detailed workflow of FeatureMAP for feature annotation and network-guided discovery of dysregulated metabolic modules.
2. **Supplementary Fig. 2.** Evidence-guided hypothesis generation, multi-agent review and final arbitration in the advanced mode of FeatureMAP.
3. **Supplementary Fig. 3.** Extended benchmarking of network-guided metabolite annotation across ten MetaboLights datasets.
4. **Supplementary Fig. 4.** Extended evaluation of FeatureMAP module identification in semi-synthetic benchmarks.
5. **Supplementary Fig. 5.** Extended evaluation of module functional interpretation using complementary embedding models and LLM-judge assessments.
6. **Supplementary Fig. 6.** Demonstration of the four core functions of PhenoMAP using purine metabolism as an example.
7. **Supplementary Fig. 7.** Community organization of phenotype-level metabolic similarity in PhenoMAP.
8. **Supplementary Fig. 8.** Supporting analyses of the Alzheimer's disease case study.
9. **Supplementary Fig. 9.** MS/MS evidence and FeatureMAP prioritization of membrane-lipid anchor metabolites in CSF and iPSCs.


#### Supplementary Tables


1. **Supplementary Table 1.** Functional capabilities of FeatureMAP and representative tools for untargeted metabolomics.
2. **Supplementary Table 2.** Scoring rubric for the forward, compatibility-oriented LLM judge.
3. **Supplementary Table 3.** Scoring rubric for the reverse, error-focused LLM judge.
4. **Supplementary Table 4.** Database information used to construct the human metabolic reaction network in FeatureMAP.
5. **Supplementary Table 5.** Curated list of ubiquitous energy and redox cofactors used for generic hub detection.
6. **Supplementary Table 6.** Software versions and key parameter settings used for method comparisons.


#### Supplementary Data


1. **Supplementary Data 1.** Candidate annotations for the MS/MS-supported cholic acid feature before and after network-guided refinement.

2. **Supplementary Data 2.** Spiked metabolites and their corresponding LC–MS features in the pathway perturbation benchmarks.
3. **Supplementary Data 3.** PathBank reference pathways used to benchmark module functional annotation.
4. **Supplementary Data 4.** Comparison of fast- and advanced-mode titles against reference pathway titles and title-rescue outcomes in the easy setting.
5. **Supplementary Data 5.** Comparison of fast- and advanced-mode titles against reference pathway titles and title-rescue outcomes in the hard setting.
6. **Supplementary Data 6.** Literature evidence and support classification for phenotype–phenotype relationships in PhenoMAP community C7.
7. **Supplementary Data 7.** Literature evidence and support classification for phenotype–phenotype relationships in PhenoMAP community C9.
8. **Supplementary Data 8.** Literature evidence and support classification for phenotype–phenotype relationships in PhenoMAP community C5.
9. **Supplementary Data 9.** Functional annotations for high-confidence host-endogenous modules ($p < 0.05$) from the clinical discovery analysis of Alzheimer's disease.
10. **Supplementary Data 10.** Functional annotations for high-confidence host-endogenous modules ($p < 0.05$) from the cellular replication analysis of Alzheimer's disease.
11. **Supplementary Data 11.** Shared and system-specific metabolite evidence for membrane-lipid remodeling in clinical discovery and cellular replication of Alzheimer's disease.
12. **Supplementary Data 12.** Shared and system-specific metabolite evidence for galactose-linked glycosphingolipid remodeling in clinical discovery and cellular replication of Alzheimer's disease.

#### Supplementary Methods

#### Supplementary Notes

1. **Supplementary Note 1**. Graph-guided exploration of purine metabolism in PhenoMAP.

#### Supplementary Figures

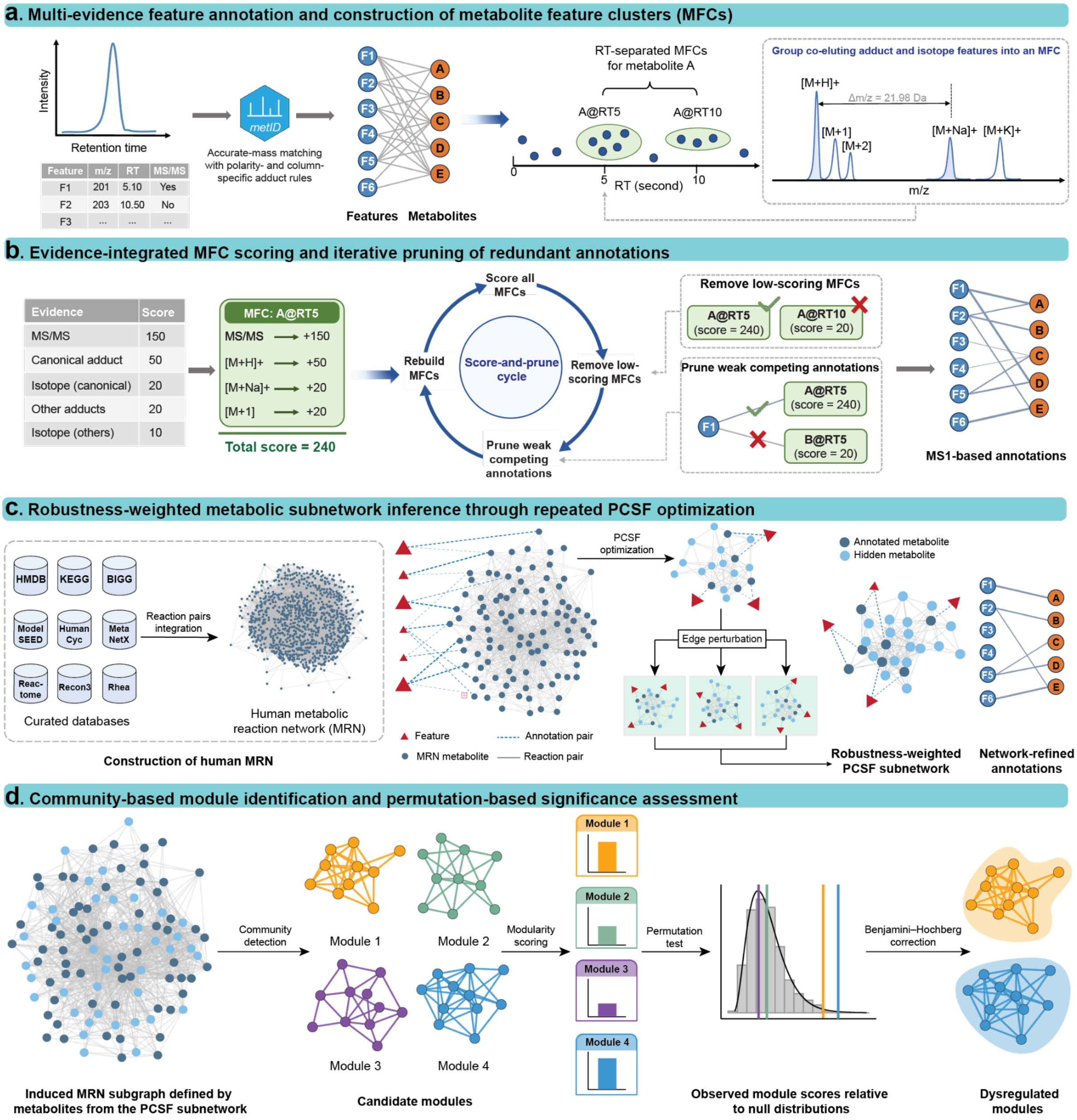


| Feature | m/z | RT | MS/MS |
|---|---|---|---|
| F1 | 201 | 5.10 | Yes |
| F2 | 203 | 10.50 | No |
| F3 | ... | ... | ... |

| Evidence | Score |
|---|---|
| MS/MS | 150 |
| Canonical adduct | 50 |
| Isotope (canonical) | 20 |
| Other adducts | 20 |
| Isotope (others) | 10 |

**Supplementary Fig. 1 | Detailed workflow of FeatureMAP for feature annotation and network-guided discovery of dysregulated metabolic modules.** (a) Multi-evidence feature-to-metabolite annotation and MFC construction. (b) Evidence-integrated MFC scoring and iterative pruning of redundant annotations. (c) Network-guided annotation refinement and robustness-weighted PCSF subnetwork construction. (d) Community-based module identification and permutation-based significance assessment.

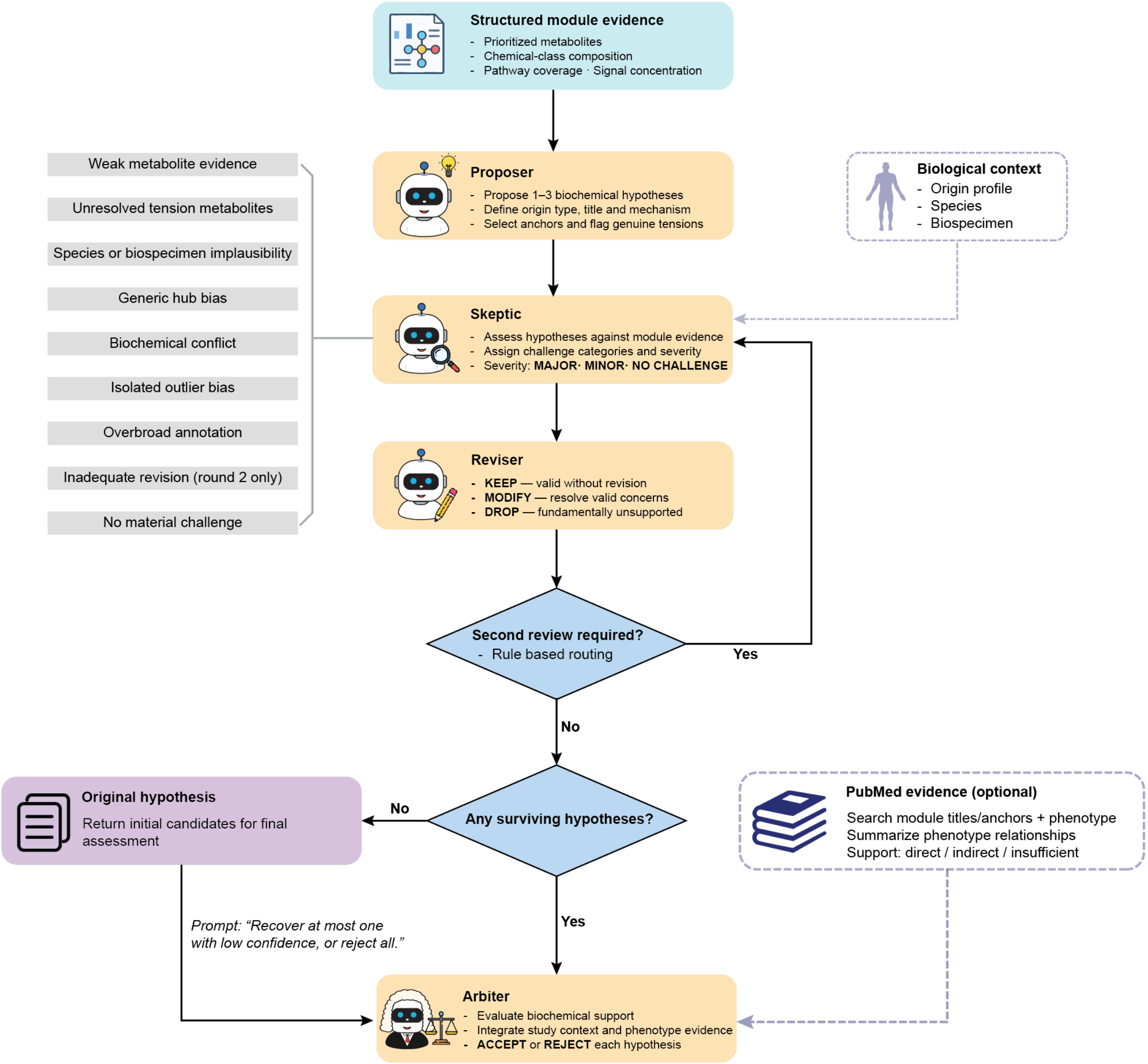


**Supplementary Fig. 2 | Evidence-guided hypothesis generation, multi-agent review and final arbitration in the advanced mode of FeatureMAP.**

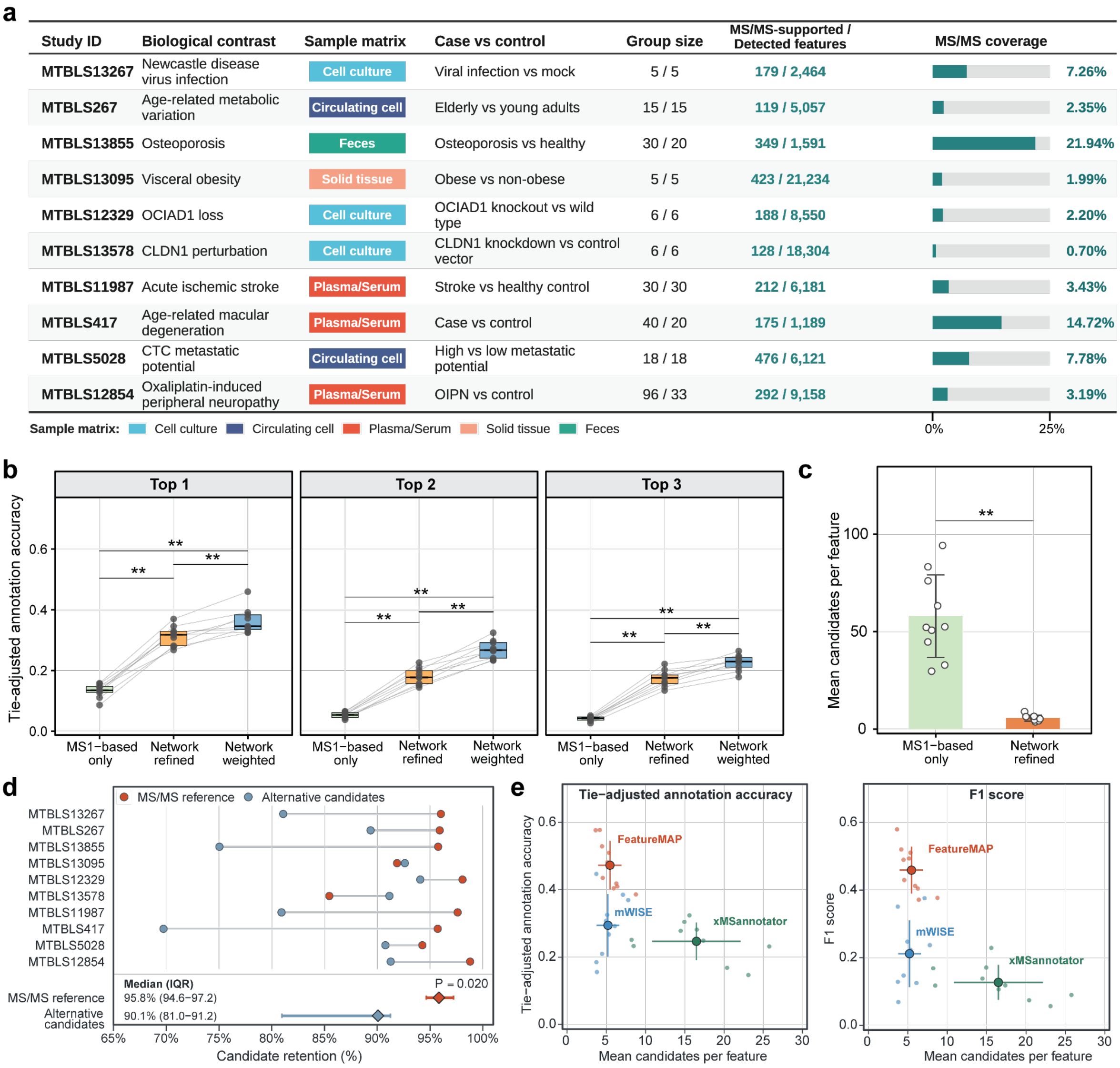


**Supplementary Fig. 3 | Extended benchmarking of network-guided metabolite annotation across ten MetaboLights datasets.** (a) Dataset characteristics and MS/MS annotation coverage. (b) Top 1-3 tie-adjusted annotation accuracy at three annotation stages. Each point represents one dataset, and lines connect measurements from the same dataset. Statistical comparisons used two-sided paired Wilcoxon signed-rank tests across ten datasets; **$P < 0.01$. (c) Mean candidate numbers per feature before and after network refinement. Each point represents one dataset. Bars and error bars indicate the across-dataset mean ± sd. Statistical comparisons used two-sided paired Wilcoxon signed-rank tests across ten datasets; **$P < 0.01$. (d) Retention of MS/MS-supported reference and alternative candidates among initial MRN-mappable annotations. Diamonds indicate across-dataset medians, and horizontal intervals show the interquartile ranges. Statistical comparisons used two-sided paired Wilcoxon signed-rank tests across ten datasets. (e) Annotation redundancy versus tie-adjusted annotation accuracy (left) and F1 score (right). Small points represent individual datasets for each tool, while large points indicate across-dataset means, with horizontal and vertical error bars showing standard deviations. Accuracy metrics were evaluated on the common MS/MS-supported feature set, whereas redundancy was calculated across each tool's full annotation output.

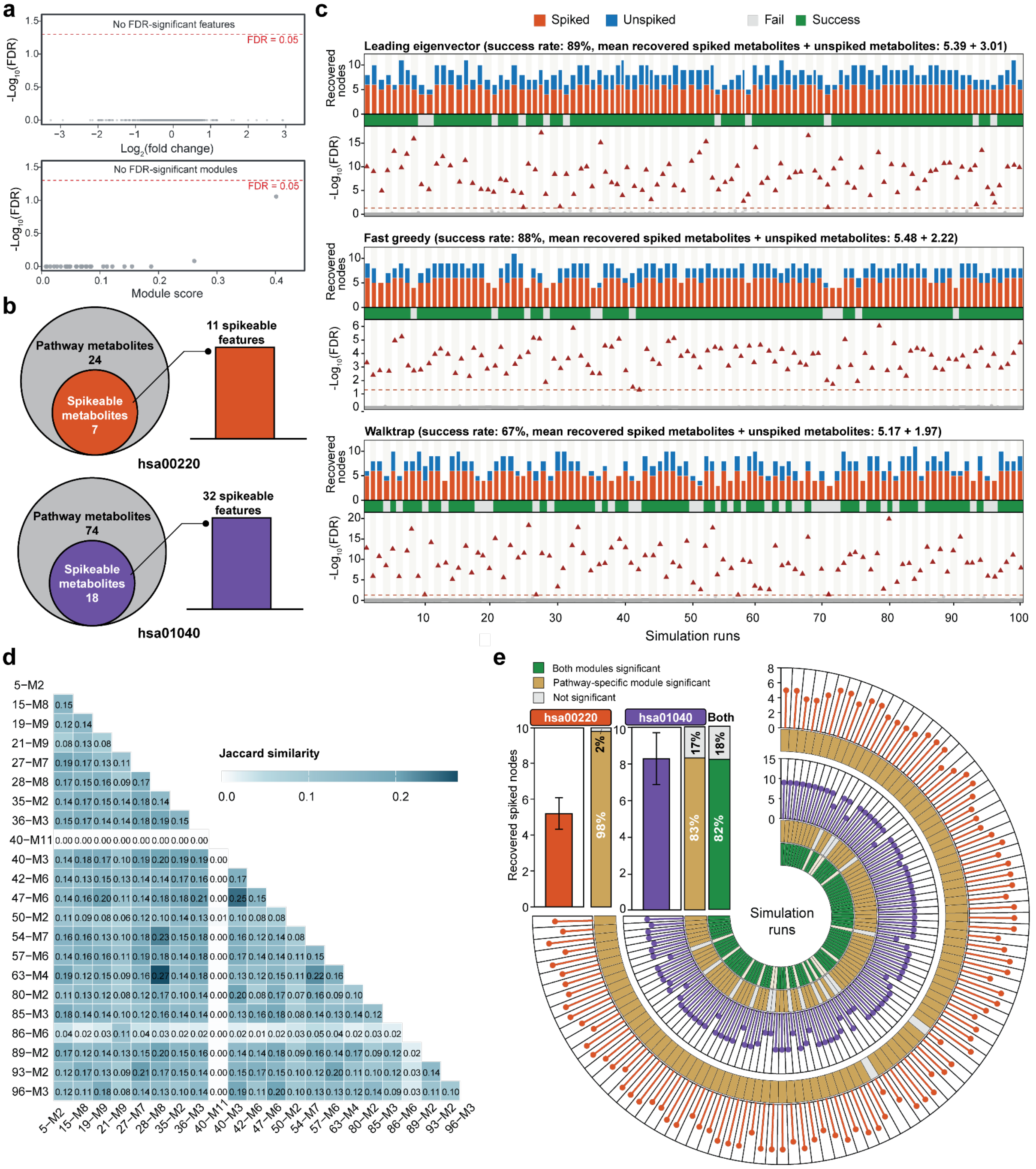


**Supplementary Fig. 4 | Extended evaluation of FeatureMAP module identification in semi-synthetic benchmarks.** (a) Baseline feature- and module-level significance before spike-in. Each point represents one feature (top) or candidate module (bottom). (b) Pathway sizes and MS/MS-supported spike-in targets for arginine biosynthesis (hsa00220) and biosynthesis of unsaturated fatty acids (hsa01040). (c) Recovery across 100 arginine-biosynthesis simulations using leading eigenvector, fast greedy and Walktrap community detection. Success required exactly one FDR-significant module containing at least five spiked metabolites. Dashed lines indicate FDR = 0.05. (d) Pairwise Jaccard similarity of metabolite sets in significant modules identified in random-metabolite negative controls. Labels indicate simulation run and module identifiers. (e) Recovery under simultaneous perturbation of both pathways across 100 simulations. Each circular sector represents one run. From outside inward, tracks show recovered spiked metabolites and selected-module significance for hsa00220 and hsa01040, followed by whether both selected modules were significant and distinct. Insets summarize mean ± s.d. of recovered spiked metabolites and the proportions of runs with significant pathway-specific modules or two significant, distinct modules.

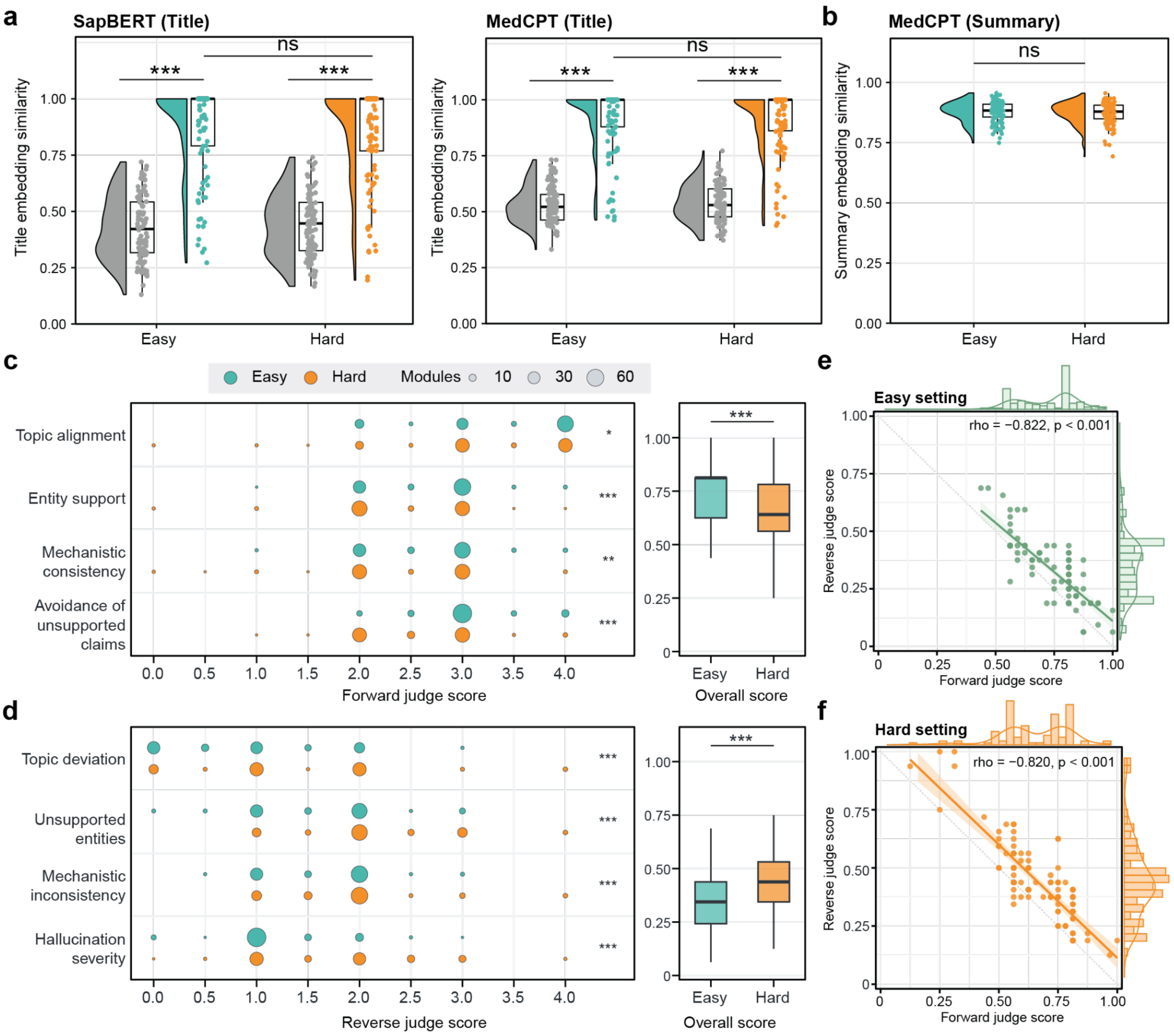


**Supplementary Fig. 5 | Extended evaluation of module functional interpretation using complementary embedding models and LLM-judge assessments.** (a) SapBERT and MedCPT embedding similarities between generated module titles and their corresponding reference titles or randomly shuffled titles. (b) MedCPT embedding similarity between generated functional summaries and reference pathway descriptions. Each point in (a,b) represents one module (n = 104 per setting). (c,d) Four-dimensional scores from the forward (c) and reverse (d) LLM judges. Bubble area represents the number of modules receiving each score. Individual dimensions retain the original 0–4 scale. Adjacent boxplots show overall scores, calculated as the mean of the four dimension-specific scores divided by four and normalized to 0–1. Higher forward scores indicate greater functional compatibility, whereas higher reverse scores indicate more severe annotation errors. Boxplots show medians and interquartile ranges. (e,f) Relationships between normalized forward and reverse overall scores in the easy (e) and hard (f) settings. Each point represents one module. Lines and shaded bands show linear fits and 95% confidence intervals. Marginal histograms and density curves show score distributions. Spearman correlation coefficients are indicated. Group comparisons in (a–d) used two-sided paired Wilcoxon signed-rank tests. *P < 0.05; **P < 0.01; ***P < 0.001; ns, not significant.

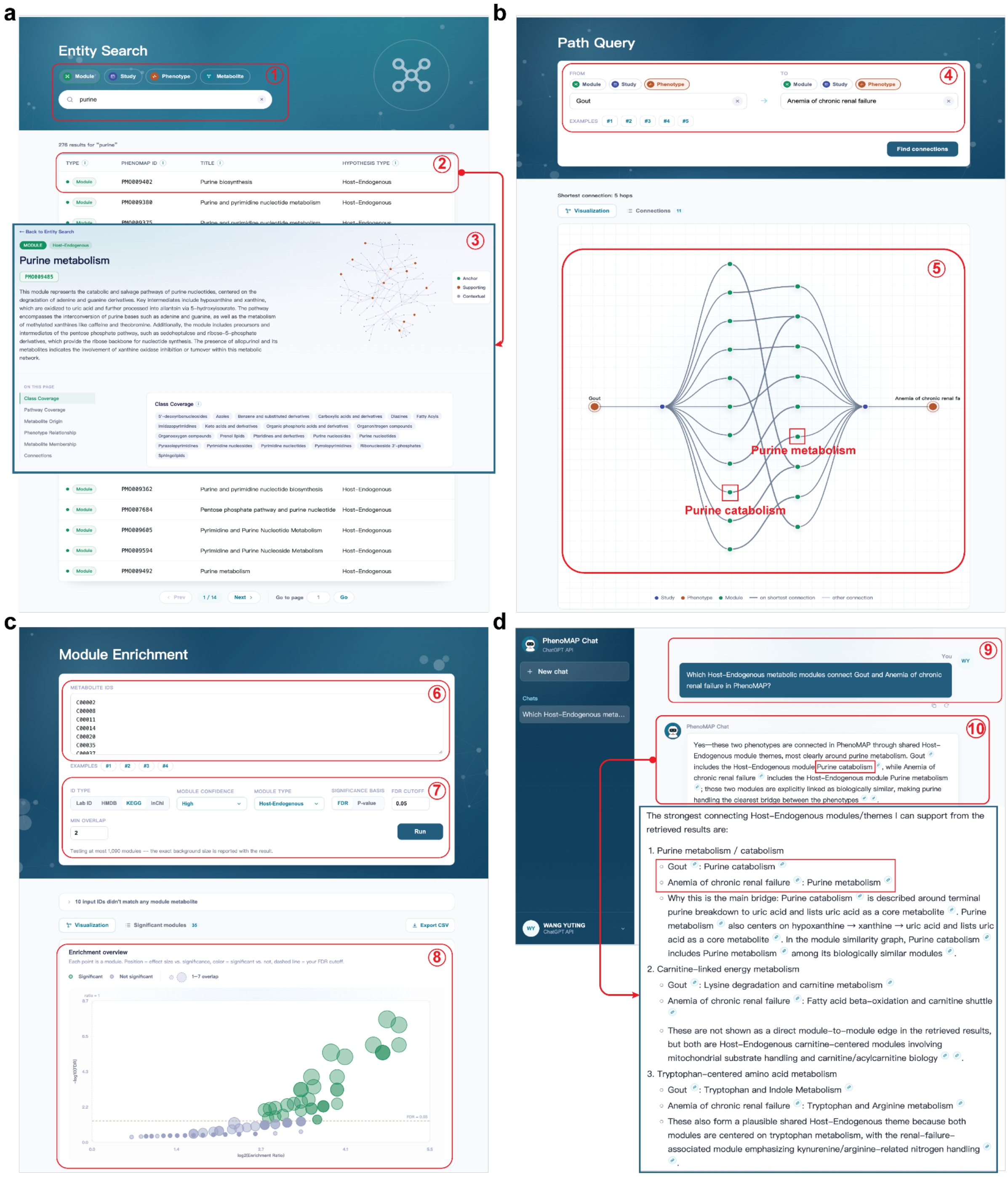


**Supplementary Fig. 6 | Demonstration of the four core functions of PhenoMAP using purine metabolism as an example.** (a) Entity Search supports entity-type selection and keyword queries (①), retrieval of matching records (②), and access to module-detail pages displaying functional annotations, metabolite networks, and supporting information (③). (b) Path Query retrieves graph connections between selected phenotypes (④). The resulting network displays graph connections between the selected phenotypes (⑤). (c) Module Enrichment accepts a metabolite list (⑥) and user-defined module filters and significance criteria (⑦). The example uses KEGG identifiers, high-confidence Host-Endogenous modules, FDR < 0.05, and a minimum overlap of two metabolites. Enrichment results are displayed as a bubble plot (⑧). (d) PhenoMAP Chat accepts a natural-language question (⑨) and returns a graph-grounded response with linked records (⑩). An enlarged response excerpt illustrates the identification of purine-related module connections and distinguishes direct graph links from broader shared metabolic themes.

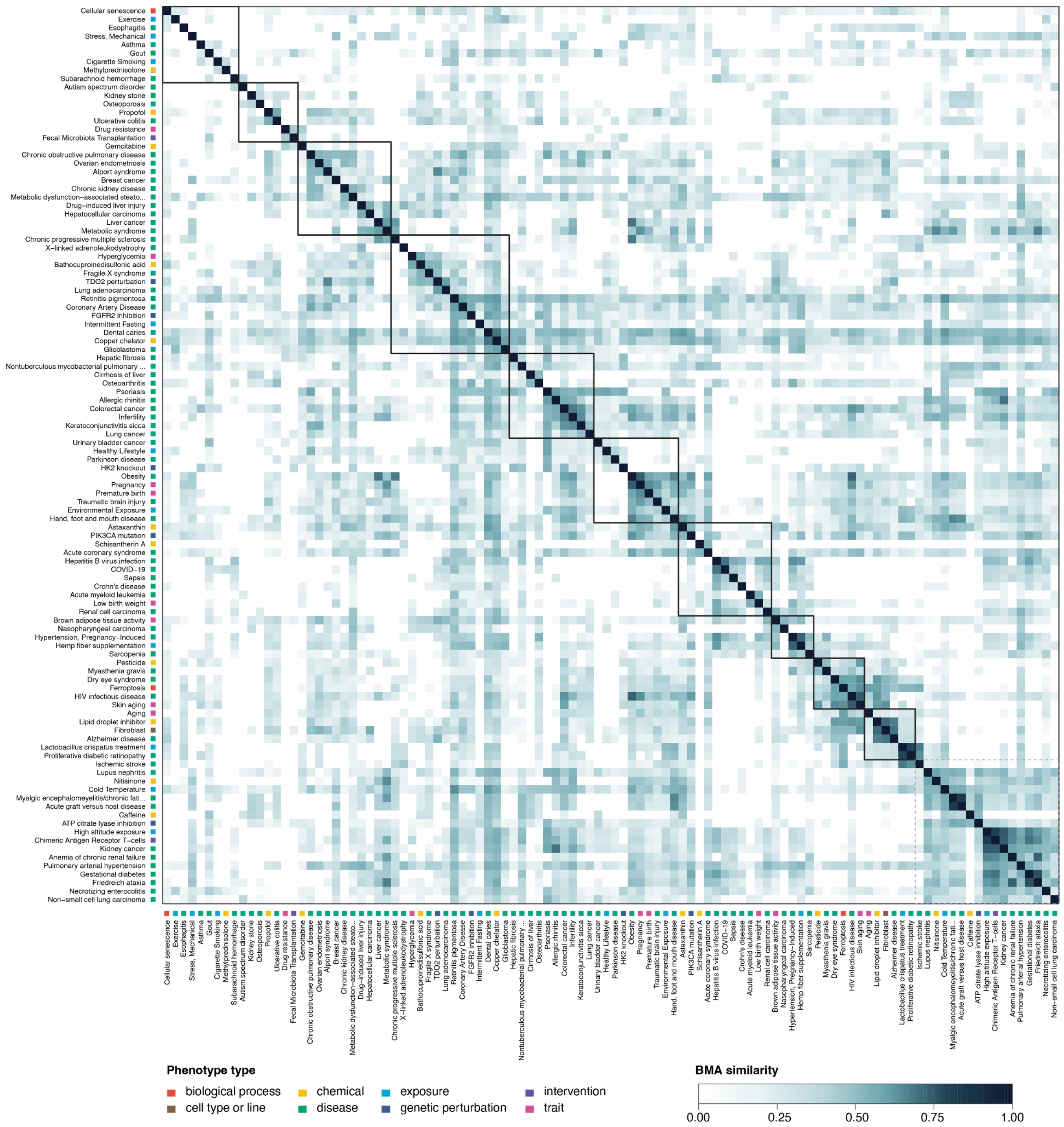


**Supplementary Fig. 7 | Community organization of phenotype-level metabolic similarity in PhenoMAP.** Heatmap of pairwise best-match-average (BMA) similarity among 106 phenotypes, each represented by at least two high-confidence, host-endogenous metabolic modules. Each cell represents the similarity between the module sets associated with two phenotypes. Phenotypes are ordered by consensus-Louvain community assignments, with communities arranged by bootstrap stability. Solid outlines delineate detected communities, whereas the dashed outline groups phenotypes without a community assignment. Colored annotation strips indicate phenotype types, and darker heatmap colors indicate greater metabolic similarity.

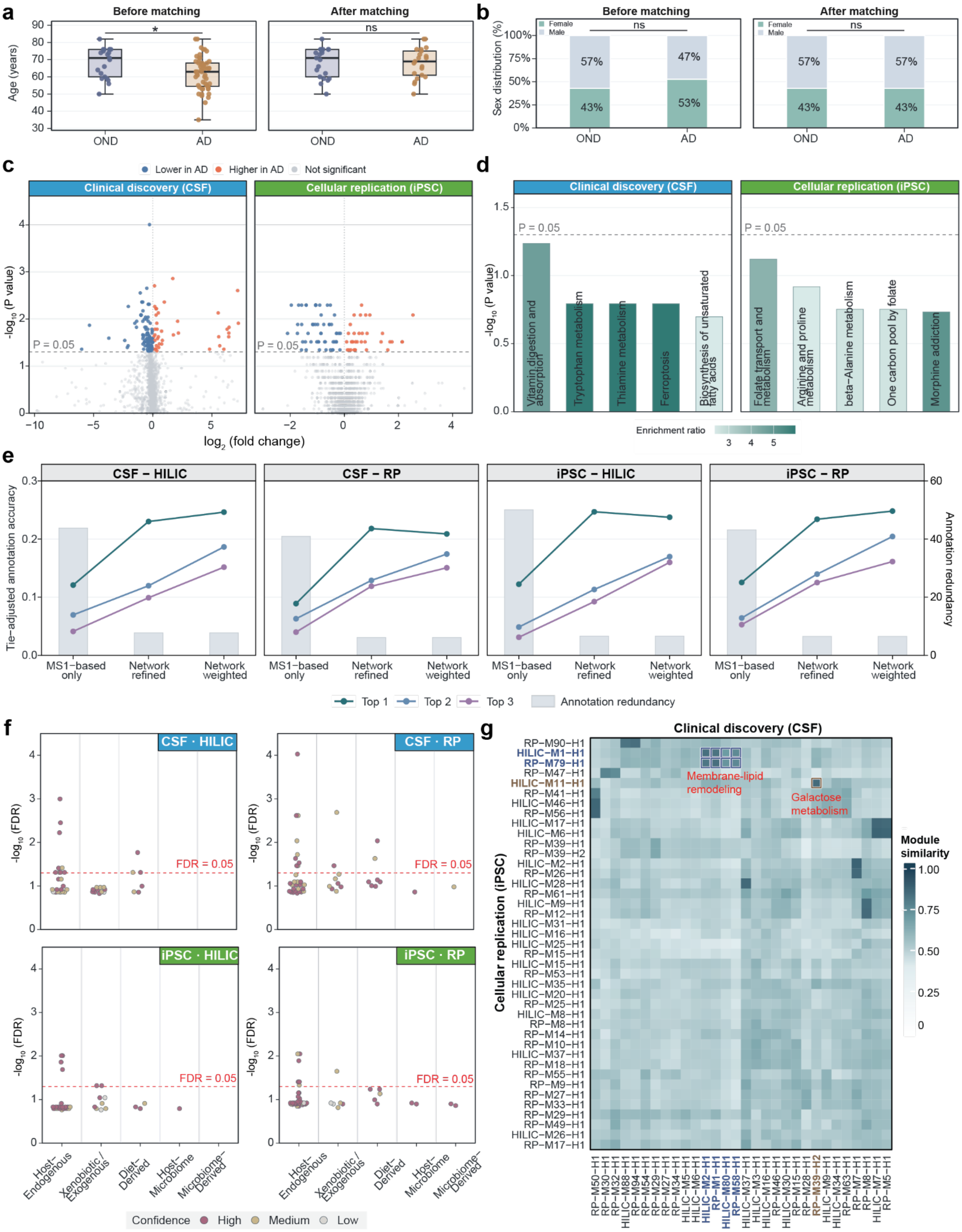


**Supplementary Fig. 8 | Supporting analyses of the Alzheimer's disease case study.** (a,b) Age and sex distributions before and after matching. Boxes show medians and interquartile ranges. Whiskers extend to 1.5 times the interquartile range. Age comparisons used two-sided Wilcoxon tests and sex comparisons used Fisher's exact tests. *P < 0.05; ns, not significant. (c) Differential analysis of Level 1/2-annotated metabolites in CSF and iPSC datasets. (d) Top five KEGG pathways ranked by enrichment P-value in each dataset. (e) Tie-adjusted annotation accuracy and candidate redundancy across annotation stages. Coloured lines show Top 1–3 accuracy against Level 1 references. Gray bars show mean candidate counts per feature. (f) Origin type, confidence, and module-level FDR of accepted hypotheses in CSF and iPSC datasets. Dashed lines indicate FDR = 0.05. (g)

Pairwise similarity between nominally significant, high-confidence Host-Endogenous module hypotheses from CSF and iPSC datasets.

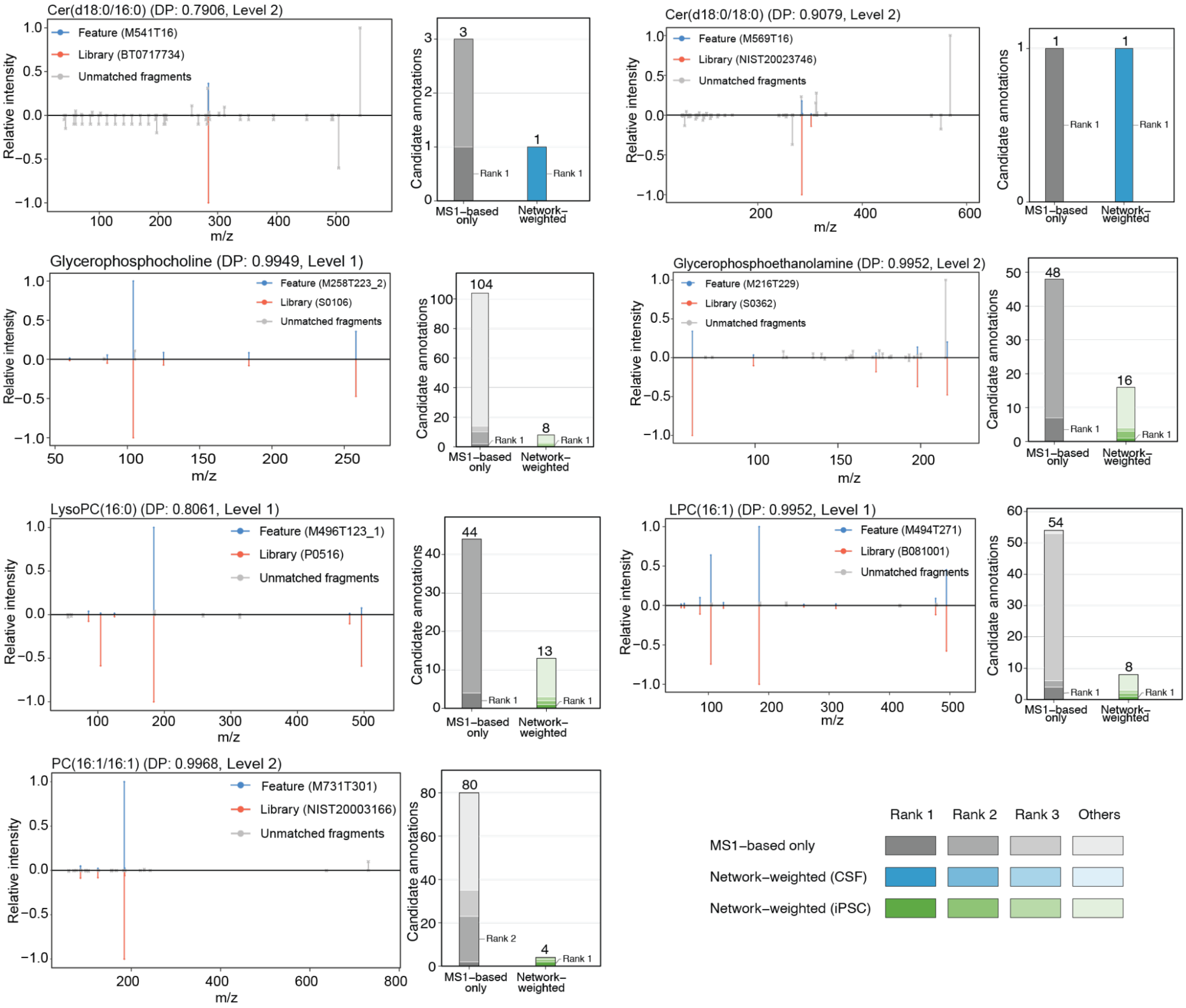


**Supplementary Fig. 9 | MS/MS evidence and FeatureMAP prioritization of membrane-lipid anchor metabolites in CSF and iPSCs.** Mirror plots compare experimental (blue) and reference (red) MS/MS spectra, with unmatched fragments shown in grey. Adjacent stacked bars show candidate annotation counts before and after FeatureMAP network weighting. Shading indicates candidate rank groups. Numbers above bars indicate total candidate counts, and labels identify the target metabolite's rank.

#### Supplementary Tables

**Supplementary Table 1. Functional capabilities of FeatureMAP and representative tools for untargeted metabolomics.**

| Category | Capability | FeatureMAP | mWISE | xMSannotator | mummichog | PIUMet |
|---|---|---|---|---|---|---|
| MS1-based annotation | Annotate LC–MS features without requiring MS/MS identification | ✓ | ✓ | ✓ | ✓ | ✓ |
| | Annotate all input features without prior filtering | ✓ | ✓ | ✓ | – | – |
| | Prioritize alternative candidates for ambiguous features | ✓ | ✓ | ✓ | ✓ | ✓ |
| | Report confidence scores for candidate annotations | ✓ | ✓ | ✓ | – | ✓ |
| Dysregulated module analysis | Identify dysregulated metabolic modules or subnetworks | ✓ | – | – | ✓ | ✓ |
| | Discover modules without prior feature filtering | ✓ | – | – | – | – |
| | Recover connections beyond predefined pathway boundaries | ✓ | – | – | ✓ | ✓ |
| | Predict unmeasured metabolites connecting observed signals | ✓ | – | – | ✓ | ✓ |
| Functional module interpretation | Module-level biological interpretation | ✓ | – | – | – | – |
| | Phenotype relevance assessment | ✓ | – | – | – | – |
| | Cross-module biological interpretation | ✓ | – | – | – | – |
| | Interpretation confidence assessment | ✓ | – | – | – | – |
| | Traceability of supporting evidence | ✓ | – | – | – | – |

**Supplementary Table 2. Scoring rubric for the forward, compatibility-oriented LLM judge.**

| Dimension | Score 4 | Score 3 | Score 2 | Score 1 | Score 0 |
|---|---|---|---|---|---|
| **Topic alignment** | Same topic, a very close pathway/sub-process, or a clearly compatible module-level topic | Mostly compatible topic with a minor scope shift or harmless generalization | Related broader/narrower metabolic topic, but less specific or incomplete | Weakly related topic with limited functional overlap | Unrelated biological function |
| **Entity support** | Key metabolites, metabolite classes, enzymes, cofactors, or biochemical entities are well supported; class-level interpretation is acceptable | Most key entities/classes are supported, with minor extras or omissions | Partial support, reasonable but incomplete class-level match, or several uncertain entities | Limited support; important entities/classes are unsupported | Mostly unsupported entities |
| **Mechanistic consistency** | Biochemical transformations, directionality, and pathway logic are compatible with the reference; detailed step-by-step coverage is not required | Mostly compatible, with minor simplification, omitted intermediates, or broader module-level logic | Plausible but broad or underspecified, or missing details important for interpretation | Mechanistic ambiguity or an unsupported mechanism that may affect interpretation | Contradicts the reference or proposes an incompatible mechanism |
| **Avoidance of unsupported claims** | No material unsupported claims; harmless background context or reasonable inference is acceptable | Minor unsupported or inferred context that does not affect interpretation | Several plausible but unverified claims or overextensions | Unsupported claims that may materially affect interpretation | Specific false, contradictory, or fabricated claims |

**Note:** Each dimension was assigned an integer score from 0 to 4. Higher scores indicate greater functional compatibility. The overall score was calculated as the mean of the four dimension-specific scores and divided by four for presentation on a 0–1 scale.

**Supplementary Table 3. Scoring rubric for the reverse, error-focused LLM judge.**

| Dimension | Score 4 | Score 3 | Score 2 | Score 1 | Score 0 |
|---|---|---|---|---|---|
| **Topic deviation** | The generated summary describes a completely unrelated biological function | The summary shifts to a different function that would materially change the interpretation | The summary is related but substantially broader or narrower in a potentially misleading way | Minor scope shift or broader pathway-family framing that remains compatible | The summary remains within the same or a clearly compatible functional topic |
| **Unsupported entities** | Central entities or entity classes are unrelated to the reference pathway | Important central entities/classes are unsupported and materially affect the interpretation | Several entities are unsupported, but the main entity class remains related | Minor additional entities, reasonable class-level additions, or plausible inferred cofactors/context | Key entities/classes are supported or compatible at the class level |
| **Mechanistic inconsistency** | Direct contradiction of the reference in directionality, intermediates, or core mechanism | Material mechanistic claim not supported by the reference and likely to change the interpretation | Mechanism is underspecified or overgeneralized in a way that could affect interpretation | Minor mechanistic imprecision, simplification, omitted intermediates, or broad module-level logic | Mechanistically consistent or compatible; detailed step-by-step coverage is not required |
| **Hallucination severity** | Multiple specific false or fabricated claims | Specific unsupported claims that materially affect interpretation | Unsupported context that could affect interpretation | Plausible extra context or reasonable inference not explicitly stated in the reference | No material unsupported claims; harmless background context is acceptable |

**Note:** Each dimension was assigned an integer score from 0 to 4. Higher scores indicate more severe annotation errors. The overall reverse-judge score was calculated as the mean of the four dimension-specific scores and divided by four for presentation on a 0–1 scale.

**Supplementary Table 4. Database information used to construct the human metabolic reaction network in FeatureMAP.**

| Name | Full name | Link |
|---|---|---|
| HMDB | Human Metabolome Database | https://www.hmdb.ca |
| KEGG | Kyoto Encyclopedia of Genes and Genomes | https://www.kegg.jp/ |
| BiGG | Biochemical, Genetic and Genomic Models | https://bigg.ucsd.edu/ |
| ModelSEED | ModelSEED Biochemistry Database | https://modelseed.org/ |
| HumanCyc | Encyclopedia of Human Genes and Metabolism | https://humancyc.org/ |
| MetaNetX | MetaNetX/MNXref metabolic network reconciliation resource | https://www.metanetx.org/ |
| Reactome | Reactome Pathway Knowledgebase | https://reactome.org/ |
| Recon3D | Three-dimensional human metabolic reconstruction | https://www.vmh.life/ |
| Rhea | Rhea knowledge base of biochemical reactions | https://www.rhea-db.org/ |

**Supplementary Table 5. Curated list of ubiquitous energy and redox cofactors used for generic hub detection.**

| Compound Name | HMDB ID | KEGG ID | Functional Category |
|---|---|---|---|
| ATP | HMDB0000538 | C00002 | Energy currency |
| ADP | HMDB0001341 | C00008 | Energy currency |
| AMP | HMDB0000045 | C00020 | Energy currency |
| GTP | HMDB0001273 | C00044 | Energy currency |
| GDP | HMDB0001201 | C00035 | Energy currency |
| Inorganic phosphate (Pi) | HMDB0001429 | C00009 | Phosphate transfer |
| Pyrophosphate (PPi) | HMDB0000250 | C00013 | Phosphate transfer |
| $NAD^+$ | HMDB0000902 | C00003 | Redox cofactor |
| NADH | HMDB0001487 | C00004 | Redox cofactor |
| $NADP^+$ | HMDB0000217 | C00006 | Redox cofactor |
| NADPH | HMDB0000221 | C00005 | Redox cofactor |
| FAD | HMDB0001248 | C00016 | Redox cofactor |
| $FADH_2$ | HMDB0001197 | C01352 | Redox cofactor |
| Coenzyme A (CoA) | HMDB0001423 | C00010 | Acyl carrier |
| $CO_2$ | HMDB0001967 | C00011 | Universal byproduct |
| $H_2O$ | HMDB0002111 | C00001 | Universal byproduct |

**Supplementary Table 6. Software versions and key parameter settings used for method comparisons.**

| Software | Version | Key parameter settings |
|---|---|---|
| FeatureMAP | 1.0.0 | ms1_match_ppm = 15; mfc_rt_tol = 5; isotope_number = 3; PCSF: b = 20, w = 25, mu = 1e-4, r = 0.2, times = 100. |
| mWISE | 0.1.0 | ppm = 10; Rt.05 = 5; method = "pearson"; diffusion.input.type = "binary"; graph.name = "RClass"; score = "z"; Unique.Annotation = TRUE. |
| xMSannotator | 1.3.2 | db_name = "HMDB"; max.mz.diff = 10; max.rt.diff = 10; cormethod = "pearson"; corthresh = 0.7; networktype = "unsigned"; deepsplit = 2; minclustsize = 10; module.merge.dissimilarity = 0.2; queryadductlist = "all". |
| mummichog | 2.7.0 | Mass tolerance: 5 ppm; feature significance cutoff: P = 0.05; permutations: 100; ion mode: positive or negative; force_primary_ion = 0. |

**Supplementary Methods**

**Raw feature annotation via evidence-integrated iterative scoring**

*Definition of evidence score*. Each MFC was assigned an evidence score by summing support from MS/MS confirmation and ion-form evidence. An MS/MS confirmation bonus of 150 points was applied once per MFC when at least one feature–metabolite pair was confirmed by MS/MS. For each ionization polarity, the presence of a canonical adduct ($[M+H]^+$ or $[M−H]^−$), isotope support for the canonical adduct, an alternative adduct, and isotope support for an alternative adduct contributed 50, 20, 20, and 10 points, respectively. Each ion-form criterion contributed at most once per polarity, regardless of the number of matching features or isotope peaks. Positive- and negative-mode evidence was accumulated within the same MFC when applicable, giving a maximum score of 350.

| Criterion present in the MFC | Score |
|---|---|
| MS/MS-confirmed feature–metabolite pair | 150 |
| [M+H]+ or [M−H]− adduct | 50 per mode |
| Isotope peak of [M+H]+ or [M−H]− | 20 per mode |
| Other adduct | 20 per mode |
| Isotope peak of another adduct | 10 per mode |

*MS/MS integration.* MS/MS annotations are optional and can be obtained independently, for example by spectral library matching in TidyMass[1]. Each candidate annotation obtained by MS1 matching is checked against the MS/MS annotations: if MS/MS assigns the same feature to the same metabolite, the candidate annotation is considered MS/MS-confirmed, and its MFC receives an additional 150 points in the evidence score. MS/MS thus increases the confidence of the MS1 candidates that it supports.

*Iterative pruning*. Each iteration consisted of three steps: (i) for each metabolite, MFCs scoring 20 or less were removed if another MFC of the same metabolite scored above 100; (ii) for each feature, candidate annotations scoring 20 or less were removed if another candidate annotation of the same feature scored above 100; (iii) the evidence scores of the remaining MFCs were recalculated. Iteration stopped when neither the mean number of MFCs per metabolite nor the mean number of candidate annotations per feature decreased further (Supplementary Fig. 1b).

**Inference of a feature-enriched metabolic reaction subnetwork via PCSF**

*Node prizes and edge costs.* In the feature–metabolite graph, node prizes and edge costs jointly determine which features, metabolites, and connections are retained by PCSF[2]. Feature nodes carry positive prizes derived from their importance scores and act as terminals that the algorithm seeks to include, whereas metabolite nodes carry non-positive prizes and are included only when they connect features at a net benefit. Metabolite prizes were defined as $\pi_m = -\mu d_m$, where $d_m$ is the degree of metabolite $m$ in the feature–metabolite graph, comprising its reaction edges in the MRN and its candidate annotation edges. The penalty parameter was set to $\mu = 10^{-4}$. At this value, the penalty is negligible for typical metabolites but becomes appreciable for highly connected cofactors. Hub metabolites are therefore selected only when they are needed to connect high-prize features, rather than serving as shortcuts between otherwise unrelated regions of the network (Supplementary Fig. 1c).

Edge costs penalize the inclusion of connections, such that weakly supported candidate annotations and long reaction paths are disfavored. For feature–metabolite edges, a constant of 0.02 was added to the cost $1 - \tilde{s}_{f,m}$ so that edges remain strictly positive in cost when $\tilde{s}_{f,m} = 1$. Metabolite–metabolite edges were assigned a uniform cost equal to the maximum feature–metabolite edge cost, capped at 0.99. Because a reaction edge is never cheaper than the least confident candidate annotation, PCSF cannot connect features more cheaply through chains of reactions than through their own annotations, and reaction edges are retained only when they link features with substantial prizes.

*Selection of β and ω.* By default, FeatureMAP uses β = 20 and ω = 25, which were used for all analyses in this study (Supplementary Table 6). Because ω exceeds the maximum prize of a single feature (β), a feature can be retained only when it is connected to other features whose combined prizes outweigh the cost of an additional tree, so that PCSF favors connected groups of dysregulated features over isolated ones. Optionally, β and ω can be tuned for a given dataset by grid search (default grid β, ω ∈ {20, 25, 30, 35, 40}), in which each combination is evaluated under 10 replicates of edge-cost perturbation and combinations are ranked by the number of features retained in the network-refined annotations, with ties broken by the mean network-weighted annotation score.

*Network-weighted annotation score.* Annotation scores frequently tie among candidate metabolites of the same feature, for example, structural isomers sharing the same adducts and isotopes, and therefore cannot rank such candidates. Edge frequencies provide complementary evidence by reflecting how consistently each candidate annotation is supported by the network across perturbed PCSF solutions. The network-weighted annotation score combines the two, with the weight α given by

$$\alpha = 0.1 + 0.2 \times \min\left(\frac{\mathrm{sd}(\varphi)}{0.25},\ 1\right)$$

where sd(φ) is the standard deviation of edge frequencies across all network-refined annotations, and 0.25, half the maximum possible standard deviation of a variable bounded in [0, 1], defines the level of dispersion at which α reaches its upper bound. When edge frequencies are similar across annotations, they carry little information for discriminating candidates, and α approaches its lower bound of 0.1. When edge frequencies vary widely, network support becomes more informative, and α increases up to 0.3. Bounding α at 0.3 ensures that annotation evidence remains the primary determinant of ranking, whereas network support mainly resolves ties and near-ties among candidates.

**Identification of functional modules via community detection**

*Construction of the induced subgraph.* Only metabolites with a node frequency above 0.05 in the PCSF subnetwork were mapped back onto the MRN, so that metabolites rarely selected across perturbed PCSF solutions did not influence module structure. The induced subgraph therefore contains all MRN reaction edges among these metabolites, including edges that were not selected in any PCSF replicate and thus have no edge frequency. Such edges were assigned the mean edge frequency of the selected edges before degree normalization.

*Metabolite-level importance.* For each annotated metabolite $m$, the metabolite-level importance $I_m$ was defined as the weighted mean of the importance scores of all features mapped to that metabolite:

$$I_m = \frac{\sum_{f \in \mathcal{F}_m} \hat{s}_{f,m} I_f}{\sum_{f \in \mathcal{F}_m} \hat{s}_{f,m}}$$

where $I_f$ denotes the importance score of feature $f$, $\hat{s}_{f,m}$ is the network-weighted annotation score of the feature–metabolite pair, and $\mathcal{F}_m$ is the set of all features mapped to metabolite $m$ in the network-refined annotations. This formulation ensures that features with stronger and more reliable annotation contribute more to the inferred metabolite importance, preventing poorly supported feature-metabolite assignments from diluting the dysregulation signal of well-characterized ones.

*Evaluation of module significance.* The statistical significance of candidate modules was assessed using a permutation-based null model. In each of $B$ permutations ($B = 100$ by default), randomized subnetworks were generated by sampling metabolites from the global MRN under the constraint that the degree-bin composition of annotated and hidden nodes matched that of the observed subnetwork used for community detection. Community detection was then applied using the same procedure as for the observed data, yielding a collection of random modules, denoted by $\mathcal{G}^{(b)}$ for permutation $b$. For each random module $H$, a module score $M_H^{(b)}$ was computed using the same definition as for the observed modules. For each permutation, importance values for randomized annotated metabolites were resampled from the observed metabolite-level importance values, so that the null model preserved the empirical distribution of dysregulation strength. The resulting collection of null module scores,

$$\mathcal{N} = \left\{ M_H^{(b)} : H \in \mathcal{G}^{(b)},\ b = 1, \dots, B \right\}$$

defined the null distribution for significance testing. A Gamma distribution was fitted to $\mathcal{N}$, and a one-sided P value was calculated for each candidate module as the probability of observing a score at least as large as the observed value under the fitted Gamma model. P values were adjusted across all candidate modules using the Benjamini-Hochberg procedure to control the false discovery rate (FDR), and candidate modules with FDR < 0.05 were designated as dysregulated modules (Supplementary Fig. 1d).

*Degree-bin-matched permutation.* Randomized subnetworks were generated on the global MRN while preserving the coarse topological composition of the observed subnetwork. Each metabolite node was assigned to a degree bin based on its degree in the full MRN, with bins defined as

$$k \leq 1, \quad 1 < k \leq 3, \quad 3 < k \leq 5, \quad 5 < k \leq 10, \quad 10 < k \leq 50, \quad k > 50$$

Within the observed subnetwork, nodes were labeled as annotated or hidden, and the numbers of annotated and hidden nodes in each degree bin were recorded as $n_A^{(j)}$ and $n_H^{(j)}$ for bin $j$, respectively. For each permutation $b$, a randomized subnetwork was constructed by sampling metabolites within each degree bin. Specifically, $n_A^{(j)}$ nodes were first sampled to form the annotated set in bin $j$, followed by $n_H^{(j)}$ nodes sampled from the remaining nodes in the same bin to form the hidden set. Sampling was performed without overlap between the two sets. When the number of available nodes was insufficient, sampling with replacement was allowed. The union of sampled nodes defined a randomized metabolite set, and the induced subgraph on the global MRN was taken as the randomized subnetwork.

**Dual-mode agentic LLM framework for functional module interpretation**

*Selection of prioritized metabolites.* To define prioritized metabolites for module interpretation, a composite priority score was calculated for each metabolite as a weighted combination of metabolite-level importance and node frequency, with weights of 0.7 and 0.3, respectively. This weighting emphasized metabolites with stronger phenotype-associated signals while still favoring those stably retained during network inference. Within each dysregulated module, metabolites were ranked in descending order by this score and assigned priority levels of high (percentile rank ≥ 0.7), medium (0.3-0.7) or low (< 0.3). The number of prioritized metabolites was determined adaptively for each module rather than fixed in advance. The relative contributions of metabolites were first modeled using Gaussian mixture modeling, and the component with the highest mean contribution was defined as the high-priority tier. When this procedure did not yield a stable partition, metabolites were instead selected from the top of the ranking until their cumulative composite priority score reached at least 20% of the module total.

*Class-level representation in the structured module context.* All metabolites within each module were summarized at the chemical-class level to provide a compact representation of the overall module composition. For each chemical class, the fraction of module metabolites assigned to that class and the names and database identifiers of member metabolites were recorded. To characterize the relative contribution of each class, the mean composite priority score across metabolites assigned to that class was calculated. Classes were then ranked within each module according to this mean score and assigned priority levels of high (percentile rank ≥ 0.7), medium (0.3-0.7), or low (< 0.3).

*Calculation of the signal concentration index.* To quantify how strongly phenotype-associated signal was concentrated within a module, a Herfindahl–Hirschman-type concentration index was calculated from metabolite-level importance values[3]. For a module $G$ with metabolite-level importance values $I_m$, the relative contribution of metabolite $m$ was defined as

$$p_m = \frac{I_m}{\sum_{m' \in G} I_{m'}}$$

The signal concentration was then calculated as

$$H_G = \sum_{m \in G} p_m^2$$

Higher values indicate that the total module signal is dominated by a small number of metabolites, whereas lower values indicate a more even distribution of signal across the module. $H_G$ was computed for all candidate modules, and each dysregulated module was assigned a high, medium, or low signal concentration category using the 70th and 30th percentiles of $H_G$ across candidate modules.

*Calculation of pathway coverage and origin profile.* To provide pathway-level biochemical context, module metabolites were mapped to curated KEGG and HMDB pathway records. For each pathway, the number of module metabolites present in the pathway and the corresponding coverage fraction, defined as this number divided by the total number of metabolites recorded for that pathway, were calculated. Pathways were ordered first by coverage fraction and then by the number of matched module metabolites, and the three highest-ranked pathways from each source were retained as compact pathway evidence for downstream LLM-based interpretation.

To provide biological-origin context, metabolite origin information was derived from HMDB records. Metabolites whose HMDB origin information contained endogenous terms were classified as endogenous, whereas those containing dietary, microbial, drug, xenobiotic, environmental, food, plant, or other exogenous source terms were classified as exogenous. The origin profile was summarized as the proportions of endogenous, exogenous, and unknown-origin metabolites.

*Prompt design for fast-mode LLM annotation*. Fast mode used a two-call prompt design to separate module-intrinsic biochemical annotation from study-context-dependent evidence evaluation. In the first call, the model was assigned the role of an expert in metabolomics and biochemical interpretation and was provided with the structured module evidence profile assembled above. This profile included the module identifier, signal concentration category, prioritized metabolites with compound names, chemical classes and priority levels, chemical-class summaries across all module metabolites, and KEGG/HMDB-derived pathway coverage. Study context variables, including species, biospecimen and phenotype, as well as optional PubMed literature evidence, were excluded from this call.

The first-call output was restricted to a JSON object with four fields: *type*, *title*, *functional_summary,* and *anchor_metabolites*. The *type* field (origin type) was constrained to one of five categories: Host-Endogenous, Microbiome-Derived, Diet-Derived, Xenobiotic/Exogenous, or Host-Microbiome. The model was instructed to assign this type according to the biological origin or dominant biochemical theme of the module. The *title* field provided a concise biochemical process name at the most specific level supported by the highest-priority metabolites. The *functional_summary* field described metabolite roles, biochemical transformations, mechanism steps, and physiological functions without referring to the study phenotype. The *anchor_metabolites* field contained several metabolites selected from the provided prioritized metabolites and represented the compounds most directly supporting the inferred interpretation.

The second-call prompt was designed to evaluate the first-call annotation in the study-specific context. The model received the outputs from the first call, together with the module origin profile, species, biospecimen, phenotype, and optional PubMed abstracts. When PubMed retrieval was enabled, literature evidence was retrieved from PubMed through the NCBI Entrez API. Separate queries were constructed from the module title and each anchor metabolite in combination with the phenotype, and up to three records were retrieved per query. The model was instructed to return a structured JSON object containing three fields: *phenotype_relationship*, *confidence_label,* and *rationale*. The *phenotype_relationship* summarized how the annotated biochemical process could relate to the study phenotype, based on PubMed evidence when available or otherwise on cautious biochemical inference. The *confidence_label* was restricted to high, medium, or low and was defined as an evidence-strength descriptor for the LLM-derived annotation. The *rationale* briefly justified the assigned confidence level by considering coherence among the module evidence, origin profile, species, biospecimen, phenotype, and available literature.

To improve parseability and consistency across modules, both calls required strict JSON output. Provider-specific response schemas were used when supported, and all responses were subjected to validation and retry-based parsing. If the first response did not contain the required fields or could not be parsed as JSON, the model was prompted again with an explicit instruction to return only the required JSON object.

*Data-layer pre-detection in advanced mode.* Before the multi-agent pipeline was initiated, each module's prioritized metabolites were screened by two rule-based checks. Generic hub detection matched compound identifiers against 16 ubiquitous energy and redox cofactors (ATP, ADP, AMP, GTP, GDP, inorganic phosphate, pyrophosphate, $NAD^+$, NADH, $NADP^+$, NADPH, FAD, $FADH_2$, coenzyme A, $CO_2$ and $H_2O$) using HMDB and KEGG identifiers (Supplementary Table 5). Isolated outlier detection flagged prioritized metabolites that were simultaneously absent from the three KEGG and HMDB pathways with the most module hits and not assigned to a high-priority chemical class of the module. When either condition was met, structured data notes naming the flagged metabolites were appended to the module evidence. These notes provided the empirical basis for the generic hub bias and isolated outlier bias challenge categories of the Skeptic.

*Agent roles and information gating.* Advanced mode was implemented as a staged peer-review workflow with role-specific information access (Supplementary Fig. 2). The Proposer generated one to three candidate biochemical hypotheses from module-intrinsic evidence only and did not assign rejection decisions. The Skeptic evaluated the proposed hypotheses using module evidence, origin profile, species and biospecimen context, but without access to phenotype or PubMed evidence. The Reviser used the same non-phenotype evidence context to respond to Skeptic critiques. Phenotype information and PubMed-derived evidence were introduced only after the internal review stage, during final arbitration by the Arbiter.

*Structured critique and revision rules.* To make the review process auditable, Skeptic critiques were constrained to nine predefined challenge categories. (1) *Weak metabolite evidence* was raised when anchor metabolites did not strongly support the claimed pathway. (2) *Unresolved tension metabolites* were raised when the hypothesis listed tension metabolites (module metabolites that the Proposer identified as difficult to reconcile with the proposed mechanism) but the summary did not explain them. (3) *Species or biospecimen implausibility* was raised when the claimed pathway was inconsistent with the stated species or sample type. (4) *Generic hub bias* was raised when the core annotation relied primarily on ubiquitous energy or redox cofactors, currency metabolites, or other non-specific intermediates. (5) *Biochemical conflict* was raised when the proposed title or mechanism conflicted with stronger pathway coverage, the dominant chemical-class composition, or several module metabolites characteristic of a different pathway. (6) *Isolated outlier bias* was raised when the core claim depended on prioritized metabolites that lacked corroboration from related metabolites, chemical class structure, pathway evidence, or biochemical plausibility. (7) *Overbroad annotation* was raised when the module evidence supported a more specific pathway, reaction family or substrate class than the proposed title reflected. (8) *Inadequate revision* was used in round 2 only, when a major challenge from round 1 had not been adequately addressed by the Reviser. (9) *No material challenge* indicated that no substantive biochemical concern was identified for a given hypothesis.

Each critique was assigned a severity label of major, minor, or no challenge. Major critiques indicated concerns that could undermine the central biochemical mechanism if unresolved, whereas minor critiques indicated limitations that warranted acknowledgment but did not invalidate the hypothesis. The Reviser's action was restricted to keep, modify, or drop. Modified hypotheses were required to update the hypothesis content. A second Skeptic-Reviser round was triggered only when a substantive modification required re-evaluation or when a major critique remained unresolved.

*PubMed retrieval and phenotype relationship summarization.* After the internal peer-review stage concluded, PubMed evidence was retrieved for surviving hypotheses through a two-layer strategy. In the first layer, anchor metabolites from all surviving hypotheses were pooled and searched collectively against the study phenotype using the NCBI Entrez API, avoiding redundant per-hypothesis API calls for shared anchors. In the second layer, each hypothesis was searched independently by title combined with the phenotype. Records from the two layers were merged per hypothesis and deduplicated by PMID, with hypothesis-specific title evidence taking priority. Retrieved abstracts for each hypothesis were then passed to a dedicated LLM summarization call that returned three structured fields: *phenotype_relationship* (2-4 sentences based solely on the retrieved abstracts, or a fixed insufficient-evidence response when no abstracts were available), *literature_support* (direct, indirect, or insufficient), and *evidence_note* (a brief characterization of evidence strength or gap). The resulting per-hypothesis summaries were supplied to the Arbiter as PubMed-derived phenotype relationship inputs.

*Arbitration and Recovery Rule.* Final verdicts were assigned only by the Arbiter. The Arbiter reviewed the surviving hypotheses together with the complete debate history, study context, origin profile, and PubMed-derived phenotype summaries, and assigned each hypothesis an *accepted* or *rejected* verdict with a confidence label and rationale. If all hypotheses had been dropped during revision, the Arbiter reviewed the original candidate set and could recover at most one biologically plausible hypothesis with low confidence. Any rescued annotation was retained only as a tentative interpretation, preventing useful but conservatively discarded hypotheses from being lost without overstating weak evidence.

*Calculation of global direction.* Global direction was calculated as a module-level output descriptor when feature-level effect sizes were available. Module metabolites were first linked to features through the network-refined annotations using their HMDB or KEGG identifiers. Each matched feature was counted once and weighted by its network-weighted annotation score $\hat{s}_f$ (the highest score if the feature was annotated to several module metabolites). The module-level direction score was calculated as the weighted mean of the effect sizes $S_f$ of the matched features:

$$D_G = \frac{\sum_{f \in \mathcal{F}_G} \hat{s}_f S_f}{\sum_{f \in \mathcal{F}_G} \hat{s}_f}$$

where $\mathcal{F}_G$ is the set of matched features of module $G$. A module was labeled as up when $D_G > 0$ and features with positive effect sizes accounted for at least two-thirds of the total weight, as down when $D_G < 0$ and features with negative effect sizes accounted for at least two-thirds of the total weight, and as mixed otherwise. Global direction was used as a statistical reference descriptor summarizing the dominant signed feature-level association within a module. It was not interpreted as a direct mechanistic direction of the underlying biochemical process and was not supplied to the LLM as a mechanistic constraint.

*Cross-module synthesis.* Cross-module synthesis was optionally performed after advanced-mode interpretation when accepted hypotheses were available from at least two modules. Each accepted hypothesis was represented as a separate node identified by its module and hypothesis identifiers. The synthesis agent received the hypothesis identifiers, origin types, functional titles and summaries, together with the study species, biospecimen and phenotype when available. The agent was instructed to group hypotheses into thematic clusters based on shared or closely related biological functions, assigning each hypothesis to at most one cluster, and to generate an overall narrative summarizing the dominant themes, cross-module connections and their potential relevance to the study phenotype.

Two complementary types of relationships were evaluated between hypotheses from different modules. Biology links were inferred by the synthesis agent from the functional titles and summaries and required a specific functional relationship, such as a shared biochemical process, substrate family or biological dependency. Each reported biology link was accompanied by a brief rationale. Biochemical links were computed deterministically, independently of the LLM. A link required both a direct MRN reaction edge connecting a prioritized metabolite from one parent module to a prioritized metabolite from the other, and at least one shared pathway identifier in the modules' KEGG or HMDB pathway-coverage records. The two evidence sources were then combined programmatically to classify each cross-module hypothesis pair as biochemical, biology, both or none. An interactive network displayed accepted hypotheses as nodes and supported cross-module relationships as edges.

**Evaluation of network-guided MS1 annotation refinement**

*Evaluated features and ranking scores.* A candidate annotation was considered correct when its HMDB or KEGG identifier matched that of the MS/MS-supported reference annotation. MS1-based and network-refined annotations were ranked by the annotation score $\tilde{s}$, and network-refined annotations were additionally ranked by the network-weighted annotation score $\hat{s}$. In the comparison with mWISE and xMSannotator, candidates were ranked by each tool's own annotation score.

*Annotation performance metrics.* For each feature, the top-$K$ set ($K \in \{1, 2, 3\}$) comprised all candidates whose scores ranked among the $K$ highest distinct score values, so that tied candidates were retained together. Tie-adjusted annotation accuracy was calculated as

$$A = \frac{1}{N} \sum_{i=1}^{N} \frac{h_i}{n_i}$$

where $N$ is the number of evaluated features, $n_i$ is the number of candidates in the top-$K$ set of feature $i$, and $h_i$ equals 1 if the correct annotation is in this set and 0 otherwise. A correct annotation therefore receives full credit only when it is ranked uniquely at the top, and partial credit when it is tied with other candidates.

For the F1 score, true positives were defined as features whose correct annotation was in the top-$K$ set, false negatives as features whose correct annotation was not, and false positives as all incorrect candidates in the top-$K$ sets, so that over-inclusive candidate lists were penalized. Annotation redundancy was calculated as the mean number of candidate annotations per annotated feature in the full annotation output.

**Spike-in simulation benchmark for module identification**

*Spike-in procedure.* Spike-in targets were features whose MS/MS annotations (top three spectral matches per feature) had KEGG identifiers belonging to the target pathway. The spiked metabolites and their corresponding LC–MS features for both target pathways are listed in Supplementary Data 2. For each target feature $j$, a fold change $FC_j$ was drawn from $U(2, 10)$, and the intensity of each pseudo-case sample was multiplied by a factor drawn from a log-normal distribution with log-mean $\log FC_j$ and log-standard deviation 0.15, introducing between-sample variability in the perturbation. Differential analysis was performed using Welch's t-test, and $\log_2$ fold changes were used as feature effect sizes. To ensure a matched comparison with mummichog, which requires a predefined list of significant features, features with P < 0.05 were used as input for both FeatureMAP and mummichog[4].

*Parameter settings.* FeatureMAP was run with $\beta = 20$, $\omega = 25$, and 100 perturbed PCSF runs. Communities were identified using the Louvain algorithm after removing metabolites with node frequency below 0.05; communities with fewer than three metabolites were discarded, and module significance was assessed with 100 permutations. To assess robustness to the choice of community detection algorithm, the single-pathway simulations were repeated using the leading eigenvector, fast greedy, and Walktrap algorithms on the same subnetworks. Mummichog (v2.7.0) was run separately in positive and negative ion modes with a mass tolerance of 5 ppm, a significance cutoff of 0.05, and 100 permutations, and modules from both modes were pooled before selecting the primary module (Supplementary Table 6).

*Two-pathway success.* For each pathway, the module containing the largest number of its pathway metabolites was selected. A run was considered successful when the selected modules for both pathways were dysregulated (FDR < 0.05) and were distinct.

**Evaluation of dual-mode LLM functional annotation**

*Pathway selection.* PathBank pathway and metabolite tables were downloaded and restricted to human pathways in the metabolic category with a curated text description, which served as the reference for evaluating functional summary[5]. MRN coverage was calculated as the fraction of HMDB identifiers in each pathway that were present among MRN nodes. When several pathways shared the same name after removal of lipid-species suffixes, only the pathway with the highest coverage was retained. The 104 retained pathways contained 4–67 MRN-mapped metabolites (Supplementary Data 3).

*Construction of semi-synthetic modules.* The semi-synthetic modules were directly formatted as FeatureMAP module-identification outputs, so that they entered functional annotation through the same procedure as modules identified from real data. Neighbor and noise metabolites were sampled without replacement from MRN nodes with HMDB identifiers, and all available nodes were used when fewer candidates than required were present within the sampling range. Each metabolite was assigned a metabolite-level importance and a node frequency, the two inputs used to select prioritized metabolites for interpretation. For pathway metabolites, both values were independently sampled from a uniform distribution between 0.80 and 1.00. For neighbor metabolites, both values were sampled between 0.60 and 0.80. For noise metabolites, both values were sampled between 0.40 and 0.60 in the easy setting, but between 0.80 and 1.00 in the hard setting. Therefore, in the hard setting, noise metabolites had the same value distribution as pathway metabolites and were equally likely to be selected as prioritized metabolites.

*Calculation of ROUGE-L recall for partially matched titles.* Before comparison, generated titles and reference pathway names were converted to lowercase and parenthetical text was removed. Title embeddings were generated using OpenAI text-embedding-3-small, SapBERT, and MedCPT, and the random control was obtained by shuffling reference names among modules of the same setting[6,7]. For lexical comparison, titles were tokenized into words at non-alphanumeric characters. A title was classified as an exact match when it was identical to the reference name, and as a partial word overlap when it shared at least one word with the reference name. For titles classified as partial word overlaps, ROUGE-L recall was defined as the length of the longest common subsequence (LCS) between the reference and generated title tokens, divided by the number of tokens in the reference title:

$$\text{ROUGE-L recall} = \frac{\text{LCS(reference title, generated title)}}{|\text{reference title}|}.$$

A higher ROUGE-L recall indicates that a larger proportion of the reference pathway title was recovered while preserving token order[8].

*Summary-level functional annotation evaluation*. Semantic similarity was quantified as the cosine similarity of text embeddings. Generated titles were compared with reference pathway names using OpenAI text-embedding-3-small, SapBERT (cambridgeltl/SapBERT-from-PubMedBERT-fulltext) and the MedCPT query encoder (ncbi/MedCPT-Query-Encoder), and functional summaries were compared with reference pathway descriptions using OpenAI text-embedding-3-large and the MedCPT query encoder. For each text pair, cosine similarity was calculated as

$$\text{Cosine similarity} = \frac{\mathbf{e}_{\text{reference}} \cdot \mathbf{e}_{\text{summary}}}{\|\mathbf{e}_{\text{reference}}\| \, \|\mathbf{e}_{\text{summary}}\|}.$$

where $\mathbf{e}_{\text{reference}}$ and $\mathbf{e}_{\text{summary}}$ denote the embedding vectors of the PathBank pathway description and the generated functional summary, respectively.

We further evaluated biological compatibility using GPT-4o as an LLM judge. The judge was provided with the reference pathway description and the generated summary and independently evaluated four dimensions: topic alignment, entity support, mechanistic consistency, and avoidance of unsupported claims. Each dimension was assigned an integer score from 0 to 4, where higher values indicated greater functional compatibility (Supplementary Table 2). The overall score was calculated as the mean of the four dimension-specific scores and divided by four for presentation on a 0–1 scale. Each summary was evaluated twice at a temperature of 0.2, and the mean score across the two evaluations was used for downstream analysis.

To reduce potential leniency in the compatibility-oriented LLM evaluation, we additionally implemented a reverse GPT-4o judge that was explicitly instructed to identify material problems in the generated summaries. Rather than assessing whether an interpretation was compatible with the reference pathway, the reverse judge evaluated the severity of topic deviation, unsupported entities, mechanistic inconsistency, and hallucinated claims on a 0-4 scale, with higher scores indicating more severe errors (Supplementary Table 3). This complementary, error-focused framing was used to examine whether the conclusions from the primary LLM judge were robust to the direction in which the evaluation question was posed.

**Construction of the PhenoMAP knowledge graph**

*Phenotype extraction and classification.* For each curated study, phenotype extraction was performed from repository-provided *study description* fields. The description text was supplied to GPT-4.1 with a constrained instruction to identify the principal biological or clinical variable defining the study comparison. The LLM was required to return a structured output containing a candidate phenotype term, one phenotype category, and a brief justification. The category definitions were specified explicitly in the prompt to reduce inconsistent classification across studies. Genetic perturbation was prioritized when the study variable was a gene knockout, knockdown, overexpression, or pharmacological perturbation of a specific gene or protein. Disease was used for human disorders or disease states, whereas symptom was reserved for clinical signs or features that did not define a full disease diagnosis. Intervention was used for procedures or therapies, chemical for named compounds or drugs, exposure for non-chemical environmental or behavioral exposures, trait for physiological or biological states, cell type/line for comparisons primarily defined by cell identity, and biological process for molecular or cellular processes. Outputs classified as unclear or not applicable were excluded from automatic ontology mapping and inspected manually.

*Phenotype-type-specific ontology mapping.* Ontology mapping was performed using phenotype-type-specific search spaces to avoid mapping the same biological concept to different ontology sources across studies. For each extracted phenotype term, candidate ontology records were retrieved from the EMBL-EBI Ontology Lookup Service only within the ontology sources assigned to its phenotype category[9]. Disease terms were searched against MONDO, EFO, MeSH and HPO; symptom terms against HPO; intervention terms against NCIT; chemical terms against ChEBI; exposure terms against EFO and MeSH; trait terms against EFO and HPO; cell type/line terms against CL and EFO; and biological process terms against GO. Genetic perturbations were not mapped through the Ontology Lookup Service; instead, the extracted gene symbol was resolved to a human NCBI Gene identifier when possible.

*Calculation of module–module similarity.* Module–module similarity was calculated to connect modules with related functional interpretations or overlapping biochemical composition across studies. For each module pair, a biological score and a compositional score were calculated. The biological score measured agreement in functional interpretation:

$$S_{\text{bio}} = 0.5\, c_{\text{summary}} + 0.4\, c_{\text{title}} + 0.1\, \delta_{\text{type}}$$

where $c_{\text{summary}}$ and $c_{\text{title}}$ are the cosine similarities of OpenAI text-embedding-3-small embeddings of the functional summaries and titles, and $\delta_{\text{type}}$ equals 1 if the two modules share the same origin type and 0 otherwise. The summary received the highest weight because it provides the most detailed description of module function. The compositional score measured overlap in module content. Metabolite overlap was first calculated as

$$S_{\text{met}} = 0.5\, J_{\text{core}} + 0.3\, J_{\text{sup}} + 0.2\, J_{\text{ctx}}$$

where $J_{\mathrm{core}}$, $J_{\mathrm{sup}}$ and $J_{\mathrm{ctx}}$ are the Jaccard indices of the core, supporting and contextual metabolites of the two modules, weighted according to their decreasing contribution to module interpretation. The compositional score was then calculated as

$$S_{\mathrm{comp}} = 0.4\,S_{\mathrm{met}} + 0.25\,J_{\mathrm{path}} + 0.25\,J_{\mathrm{class}} + 0.1\,\delta_{\mathrm{type}}$$

where $J_{\mathrm{path}}$ and $J_{\mathrm{class}}$ are the Jaccard indices of the KEGG and HMDB pathways and of the chemical classes assigned to the two modules. The two scores were combined as

$$S = 0.75\max(S_{\mathrm{bio}}, S_{\mathrm{comp}}) + 0.25 \cdot \frac{S_{\mathrm{bio}} + S_{\mathrm{comp}}}{2}$$

so that module pairs strongly supported by either functional interpretation or biochemical composition were retained, while pairs supported by both received the highest scores. Module pairs with $S \geq 0.70$ were connected by "similar to" relationships and labeled as high ($S \geq 0.75$) or medium ($0.70 \leq S < 0.75$) similarity.

*Module enrichment analysis.* Input metabolites could be provided as PhenoMAP, HMDB or KEGG identifiers or InChI strings. They were mapped to PhenoMAP metabolite nodes using the selected identifier type, and unmatched identifiers were reported separately. Each module was represented by a module set comprising its core and supporting metabolites. The background was defined as the union of all module sets, restricted to metabolites with an identifier of the selected type, because only these metabolites could be matched by the input. Input metabolites outside the background were not tested.

For each module, let $N$ be the background size, $K$ the module set size, $n$ the number of input metabolites in the background and $k$ the number of input metabolites in the module set. Over-representation was assessed using a one-sided hypergeometric test:

$$P = \sum_{i=k}^{\min(K,n)} \frac{\binom{K}{i}\binom{N-K}{n-i}}{\binom{N}{n}}$$

and the enrichment ratio was calculated as $(k/n)/(K/N)$. Benjamini–Hochberg correction was applied across modules with $k \geq 1$, because modules sharing no metabolite with the input cannot show enrichment. By default, modules with FDR < 0.05 and $k \geq 2$ were reported as enriched. Users can optionally restrict the analysis to modules with selected confidence labels or origin types, in which case the background was rebuilt from the retained modules. For each module, the overlap count, enrichment ratio, nominal P value, FDR, and overlapping metabolites were reported.

*Natural-language query routing and tool-grounded response generation.* PhenoMAP Chat processed each user message in three stages. In the first stage, the message was classified as a question about

PhenoMAP graph data, a question about the PhenoMAP resource itself (for example, its functions or how to cite it), or an unrelated request. Resource questions were answered using a dedicated tool that retrieves curated product information, and unrelated requests were declined with a brief explanation. In the second stage, graph-data questions were assigned to one of six intents, and the entities mentioned in the question were extracted: lookup (finding an entity), profile (retrieving the details of one entity), association (retrieving the graph neighbors of one entity), connection (finding how two entities are related), enrichment (testing a metabolite list), and anchored query (a question involving a module named by its title). No graph records were retrieved in the first two stages.

In the third stage, the chat service called the PhenoMAP tools corresponding to the web interface functions: entity search, node-detail retrieval, path query, and module enrichment. When a user-provided name was resolved to an entity with a different name, the user was asked to confirm the match before an answer was generated. The final response was generated only from the records returned by these tools in the current turn. Each retrieved study, phenotype, module and metabolite was recorded in a citation dictionary keyed by its PhenoMAP identifier, and identifiers cited in the response were rendered as links only when they were present in this dictionary, so that every cited statement could be traced back to the graph records used to generate it.

*Community study submission and curation.* The PhenoMAP Community page accepts user-submitted metabolomics studies for review and future graph expansion. Each submission includes study metadata, including title, phenotype, optional phenotype identifier, study design, sample size, biospecimen, MS/MS availability, chromatography, mass spectrometry, study summary, DOI or source link and submitter contact information. Users are required to provide a feature table and a sample metadata file, and may optionally provide a metabolite annotation file. Each submission is assigned a submission identifier and stored separately from the public PhenoMAP graph. Submitted studies are not automatically incorporated into PhenoMAP. They are reviewed for metadata completeness, study design, data suitability and compatibility with the FeatureMAP workflow, and accepted studies are processed using the same phenotype-normalization, FeatureMAP analysis and graph-construction procedures as repository-derived studies before inclusion in a future release.

**Cross-study analyses in PhenoMAP**

*Module selection and BMA similarity.* Cross-study analyses used modules with a high confidence label and the Host-Endogenous origin type, comprising 780 modules from 158 phenotypes. For two module sets $A$ and $B$, BMA similarity was calculated as

$$\mathrm{BMA}(A,B)=\frac{1}{2}\left(\frac{1}{|A|}\sum_{a\in A}\max_{b\in B}S_{ab}+\frac{1}{|B|}\sum_{b\in B}\max_{a\in A}S_{ab}\right)$$

where $S_{ab}$ is the module–module similarity between modules $a$ and $b$[10]. Phenotype-level similarity was calculated for the 106 phenotypes represented by at least two modules.

*Phenotype community detection.* A phenotype network was constructed by connecting phenotype pairs whose BMA similarity ranked within the top 5% of all pairs. Louvain clustering was performed with a resolution parameter of 2.0. Because Louvain clustering depends on the order in which nodes are processed, a consensus approach was used. Louvain clustering was run 50 times with randomly shuffled node order, and for each pair of phenotypes, the fraction of runs in which they were assigned to the same community was recorded. Phenotype pairs grouped together in at least half of the runs were connected in a consensus network, and Louvain clustering of this network defined the final communities. Communities with fewer than three phenotypes were not considered. The stability of each community was assessed by bootstrap resampling: in each of 50 resamplings, 85% of phenotypes were randomly selected and clustered again using the same procedure, and the Jaccard similarity between the original community and its most overlapping community in the resampled result was calculated. The average across resamplings was used as the community stability, with values close to 1 indicating that the community was recovered almost unchanged.

*Literature curation of phenotype connections.* Each connected phenotype pair was searched in the literature, and the evidence was recorded as one or more of five types: population or clinical studies linking both phenotypes (HUM); experiments in which one phenotype was manipulated and the other was affected (EXP); described molecular, biochemical or regulatory relationships between the two (MECH); independent links of both phenotypes to the same specific mechanism (SHARED); and ontological, compositional or definitional relationships (STRUCT). Pairs supported by HUM, EXP, MECH, or STRUCT evidence were classified as directly supported, and pairs supported only by SHARED evidence as indirectly supported. Broad mechanisms, such as inflammation, oxidative stress, or general mitochondrial dysfunction, were not considered sufficient for indirect support. Curated evidence and support classifications for communities C7, C9 and C5 are provided in Supplementary Data 6-8, respectively.

**Supplementary Note 1. Graph-guided exploration of purine metabolism in PhenoMAP.**

We used purine metabolism to illustrate how PhenoMAP can connect a biochemical theme to the modules, phenotypes and studies represented in the knowledge graph. An Entity Search for "purine" retrieved modules with purine-related functional annotations. Individual results could then be opened as module-detail records containing the functional interpretation, metabolite network, pathway and chemical-class coverage, module origin and links to the supporting study and phenotype (Supplementary Fig. 6a). This provides an entry point for examining how a broadly defined metabolic process is represented across independently analyzed metabolomics studies.

We next used Path Query to examine the graph connections between gout and anemia of chronic renal failure. The returned paths included a Host-Endogenous module annotated as "Purine catabolism" in gout and a related module annotated as "Purine metabolism" in anemia of chronic renal failure (Supplementary Fig. 6b). These modules were connected by a module–module similarity relationship, providing a traceable metabolic bridge between the two phenotypes even though the contributing studies did not produce identically named modules. This illustrates how PhenoMAP uses similarity in module composition and functional interpretation to identify related metabolic signals across heterogeneous studies.

Module Enrichment provided a complementary route for entering the atlas from an external metabolite list. Metabolites supplied as KEGG identifiers were mapped to PhenoMAP metabolite records and tested for over-representation in the core and supporting metabolite sets of high-confidence Host-Endogenous modules. Significant modules were returned together with their overlap counts, enrichment ratios and statistical significance (Supplementary Fig. 6c). This function allows metabolite lists generated outside the FeatureMAP workflow, including results from annotated untargeted or targeted assays, to be compared with metabolic modules collected across PhenoMAP without repeating feature-level analysis.

Finally, we asked PhenoMAP Chat which Host-Endogenous metabolic modules connected gout and anemia of chronic renal failure. The query was resolved to the corresponding phenotype records and routed to the graph-search functions. The response identified purine catabolism and purine metabolism as the clearest graph-supported bridge and linked the relevant records, while distinguishing this direct module-level connection from broader metabolic themes that were shared between the phenotypes but were not represented by an explicit graph path (Supplementary Fig. 6d). Thus, PhenoMAP Chat provides a natural-language layer for interrogating the atlas while retaining traceability to the studies, modules and relationships used to generate the response.
